# ETSAM: Effectively Segmenting Cell Membranes in cryo-Electron Tomograms

**DOI:** 10.1101/2025.11.23.689996

**Authors:** Joel Selvaraj, Jianlin Cheng

## Abstract

Cryogenic Electron Tomography (cryo-ET) is an emerging experimental technique to visualize cell structures and macromolecules in their native cellular environment. Accurate segmentation of cell structures in cryo-ET tomograms, such as cell membranes, is crucial to advance our understanding of cellular organization and function. However, several inherent limitations in cryo-ET tomograms, including the very low signal-to-noise ratio, missing wedge artifacts from limited tilt angles, and other noise artifacts, collectively hinder the reliable identification and delineation of these structures. In this study, we introduce ETSAM - a two-stage Segment Anything Model 2 (SAM2)-based fine-tuned AI method that effectively segments cell membranes in cryo-ET tomograms. It is trained on a diverse dataset comprising 83 experimental tomograms from the CryoET Data Portal (CDP) database and 28 simulated tomograms generated using PolNet. ETSAM achieves state-of-the-art performance on an independent test set comprising 10 experimental tomograms for which ground-truth annotations are available. It robustly segments cell membranes with high sensitivity and precision, significantly outperforming existing deep learning methods.

## 1 Introduction

Cryo-electron tomography (cryo-ET) has emerged as a powerful imaging technique for visualizing the three-dimensional (3D) architecture of biological specimens at near-atomic resolution in their native cellular environment. In cryo-ET, samples such as cells or tissues are rapidly frozen to preserve their native state. The frozen specimen is then imaged using a transmission electron microscope, where a series of two-dimensional (2D) projections is acquired by tilting the sample over a range of angles, typically ±60°. These projections are computationally reconstructed into a 3D volume [1–3]known as a tomogram, which provides detailed insights into subcellular structures, macromolecular assemblies, and dynamic cellular processes.

Cell membranes serve as dynamic barriers that compartmentalize cellular activities, facilitate signaling, and mediate interactions with the extracellular environment. Membranes play a vital role in many cellular functions such as vesicle trafficking [4, 5], synaptic transmission, conversion of energy in chloroplast thylakoids [6–10] and mitochondrial cristae [11–16], viral entry and budding [17–24], cell division [25, 26], organelle biogenesis [27], and the localization of membrane-associated proteins [28]. Segmenting membrane structures in cryo-ET tomograms enables researchers to identify membrane morphology [29] and curvature [30], thereby facilitating understanding of cellular function. For instance, precise membrane segmentation can help reveal how viruses like HIV [31, 32] or SARS-CoV-2 [19, 20] exploit host membranes during infection, and how defects in membrane architecture contribute to neurodegenerative diseases like Alzheimer’s disease [33, 34]. Therefore, accurate membrane segmentation in cryo-ET tomograms has emerged as an important task in computational biology [35].

Traditional methods for membrane segmentation rely on manual or semi-automated approaches, often incorporating image processing techniques to enhance and delineate membrane features. One prominent example is TomoSegMemTV [36], which uses the scale-space operation based on Gaussian filtering to isolate features at a given membrane thickness, followed by the detection of ridge-like membrane regions via a Hessian tensor or edge-like membrane regions via a structure tensor. Further, it employs a tensor voting mechanism that enables voxels within the same membrane to reinforce one another, fill gaps, and disregard attached structures. However, it requires human intervention and parameter tuning to segment cell membranes from cryo-ET tomograms. Moreover, traditional approaches struggle to account for variability in membrane appearance across different imaging conditions.

In recent years, deep learning methods have gained traction as promising alternatives for automating membrane segmentation in cryo-ET. Techniques such as Membrain-Seg [37] employ a U-Net-based model trained on annotated tomograms to predict membranes. It leverages the learned features to handle noise and variability more robustly than traditional methods. Similarly, TARDIS (Transformer-based Rapid Dimensionless Instance Segmentation) [38] uses a custom-designed Convolutional Neural Network (CNN) called FNet to identify and delineate cell membranes. TARDIS was trained on diverse annotated (labeled) tomograms. It also integrates a point-cloud-based DIST (Dimensionless Instance Segmentation Transformer) model for instance segmentation of predicted cell membranes. These deep learning-based approaches have demonstrated superior performance on benchmarks, reducing segmentation time from hours to minutes and enabling the scalable analysis of large cryo-ET datasets. However, membrane segmentation remains an open problem, as current deep learning models fail to achieve both high precision and recall without tradeoffs. Some models often miss critical membrane regions in their predictions due to low sensitivity, while other more sensitive models are prone to noisy predictions.

The challenges facing membrane segmentation in cryo-ET tomograms are largely due to their inherent limitations [39–41]. Tomograms often suffer from a low signal-to-noise ratio (SNR) because electron doses must be minimized to prevent radiation damage to biological samples, resulting in noisy 3D reconstructions in which membrane boundaries are faint and difficult to distinguish from surrounding cytosolic or extra-cellular densities. Additionally, the missing wedge problem [42] arises from the limited tilt range during data acquisition, leading to incomplete sampling in the Fourier space and anisotropic resolution in the reconstructed tomogram. This manifests as elongation or blurring along the direction of the electron beam, distorting membrane shapes and complicating the identification of curved or closely apposed structures. Furthermore, sample preparation artifacts, such as ice contamination or uneven vitrification, introduce background scattering that diminishes membrane contrast and often creates crystalline patches or blobs that resemble membrane-like edges, leading to potential false positives in segmentation.

In this study, we introduce ETSAM, a fine-tuned two-stage Segment Anything Model 2 (SAM2)-based [43] AI model for effectively segmenting cell membranes in cryo-ET tomograms. SAM2 is a foundational image and video segmentation model that can accurately segment objects in natural images and videos but cannot be applied to cryo-ET tomograms directly. By repurposing SAM2’s video segmentation pipeline, ETSAM treats consecutive tomogram slices as a sequence of video frames. ETSAM leverages SAM2’s memory encoder and memory attention mechanisms to effectively identify and track membranes across each slice of the tomogram. This approach enables the model to maintain consistency when tracking complex, curved membrane structures while mitigating some inherent limitations of cryo-ET, such as low signal-to-noise ratio and missing wedge artifacts.

To train ETSAM, we collected 83 experimental tomograms from the CryoET Data Portal (CDP) [44] database along with their corresponding membrane annotations, and supplemented them with 28 simulated tomograms generated using PolNet [45]. Since most membrane annotations from the CDP database are generated using AI tools with minimal human intervention, they often contain noise artifacts, false-positive predictions, or missing membrane regions. The selected 83 experimental tomograms are among the ones with minimal missing membrane regions in the CDP database. To further improve the quality, we reduced noise by manually cleaning the annotations. The first stage of ETSAM was extensively trained on all 111 tomograms in the training dataset. The fine-tuned first-stage ETSAM was then applied back to the training dataset to generate the membrane segmentation. The first-stage predicted membrane segmentation and the original tomograms of the training data were fused together as input to train the second stage of ETSAM, which outputs the final segmentation.

We created an independent test set comprising 10 experimental tomograms, for which we manually annotated the ground truth using existing annotations from the CDP database and AI-predicted annotations as a starting point, then manually cleaned and corrected them. ETSAM achieved state-of-the-art results on the experimental test dataset compared to existing deep learning-based methods by attaining the highest Dice and Intersection over Union (IoU) scores. Moreover, ETSAM achieves the highest Area Under the Precision-Recall Curve (AUPRC) scores, indicating it outperforms existing methods across a broad range of cutoff thresholds and thereby offers the best overall precision-recall trade-off.

## 2 Results

To assess the quality of membrane segmentations, we use various evaluation metrics such as Dice, IoU, precision, recall, and AUPRC, and report the average scores across the test tomograms. Precision measures the accuracy of the model’s positive membrane predictions, whereas recall measures its completeness. Dice score, also known as F1 score, is the harmonic mean of precision and recall, providing a single metric that balances them equally. The IoU score is the ratio of the area of intersection between the predicted segmentation and the ground truth to the area of their union, measuring the extent to which they overlap. The AUPRC score measures the area under the Precision-Recall (PR) curve, plotted at various probability thresholds of the final predictions. ETSAM is compared with two other leading deep learning-based membrane segmentation methods: Membrain-Seg [37] and TARDIS [38]. The default parameters are used for both methods. To evaluate the raw performance of each method, post-processing and test-time augmentation were not applied.

### 2.1 Evaluation on Experimental Tomograms

ETSAM was evaluated on 10 experimental test tomograms from 8 CDP datasets of 5 different cellular organisms and 1 virus. As shown in Table 1, ETSAM delivers superior performance compared to Membrain-Seg and TARDIS across all metrics. The highest Dice (0.7867) and IoU (0.6751) scores indicate greater overlap between the predicted segmentation and the ground-truth membrane regions. Additionally, it achieves the highest recall (0.8025) and precision (0.8016), indicating that it captures the highest proportion of membrane regions with the best precision. Membrain-Seg has the second-highest recall of 0.7362, but its precision is significantly lower at 0.4741. TARDIS’s precision of 0.6202 is higher than Membrain-Seg’s, but its recall is substantially lower at 0.5742. Figure 1a and Supplementary Table 1 show the per-tomogram comparison of ETSAM, Membrain-Seg, and TARDIS. It can be observed that ETSAM consistently achieves better Dice and IoU scores across most experimental tomograms.

**Table 1.** The comparison of ETSAM and two other methods on 10 experimental tomograms from the test dataset in terms of multiple evaluation metrics.

| Method | Dice/F1 | IoU | Precision | Recall | AUPRC |
| --- | --- | --- | --- | --- | --- |
| ETSAM | **0.7867** | **0.6751** | **0.8016** | **0.8025** | **0.8209** |
| Membrain-Seg | 0.5399 | 0.4103 | 0.4741 | <u>0.7362</u> | 0.5351 |
| TARDIS | <u>0.5553</u> | <u>0.4111</u> | <u>0.6202</u> | 0.5742 | <u>0.5928</u> |

Bold denotes the best result, and underline denotes the second best result.

**Fig. 1.**
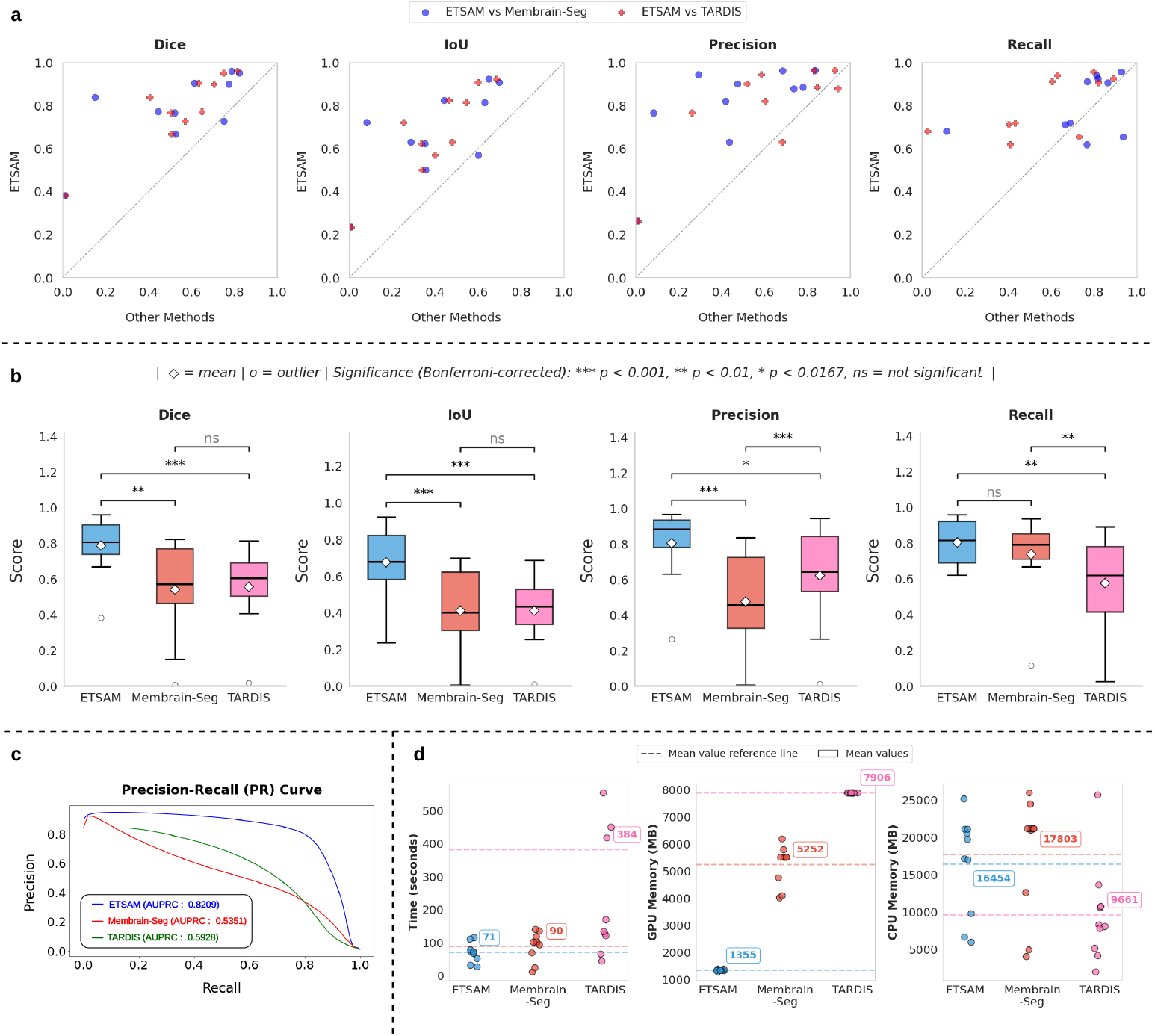
Comparison of ETSAM, Membrain-Seg, and TARDIS across various evaluation metrics on 10 experimental tomograms from the test dataset. **(a)** Parity plots that show per-tomogram performance of ETSAM against Membrain-Seg and TARDIS in terms of multiple segmentation metrics. Points above the dotted diagonal line indicate higher scores for ETSAM than the other methods. **(b)** Box plots to compare the distribution of each metric’s scores attained by all three methods, along with the statistical significance markers based on the p-values attained from a Bonferroni-corrected paired Student t-test. **(c)** Compares the Precision-Recall (PR) curve for all three methods along with the computed AUPRC scores. **(d)** Benchmark of runtime, CPU, and GPU memory compute usage of all three methods.

To analyze the precision vs recall trade-off at different thresholds, we plotted the average Precision-Recall (PR) curve in Figure 1c and computed the AUPRC for ETSAM, Membrain-Seg, and TARDIS-generated membrane segmentation. We used the probability scores predicted by each method for every voxel across all tomograms in our test set to compute the AUPRC score. It can be observed that ETSAM achieved the best AUPRC score of 0.8209 compared to TARDIS’s 0.5928 and Membrain-Seg’s 0.5351. This highlights ETSAM’s best precision-recall trade-off and its ability to achieve high recall scores while maintaining state-of-the-art precision. It indicates that ETSAM can accurately predict more membrane regions while maintaining a significantly low false-positive rate.

To analyze the significance of ETSAM results over Membrain-Seg and TARDIS, we computed a multi-variable paired Student’s t-test with Bonferroni correction on the following evaluation metrics: Dice, IoU, Precision, and Recall score. Prior to computing the t-test, we performed the Shapiro-Wilk test and ensured that the normality of differences assumption is satisfied for every paired t-test. In Figure 1b, we have used a box plot to compare the distribution of each metric across all three methods, along with the significance markers based on the p-value attained from the t-test. For comparing three methods with a given significance level of 0.05, we use the Bonferroni corrected *α* = 0.05/3 = 0.0167 as the condition to reject the null hypothesis. If the computed *pvalue > α*, there is no statistically significant difference between the pair of methods compared for the given metrics, and it is denoted by “*ns*” in Figure 1b. If *pvalue* < *α*, it is considered statistically significant and denoted with “∗”. If *pvalue* < 0.01, it is denoted with “∗∗”, and if *pvalue* < 0.001, it is denoted with “∗∗∗” to indicate higher significances. It can be observed that ETSAM shows statistical significance on Dice, IoU, and precision scores against both Membrain-Seg and TARDIS. For the recall score, ETSAM shows statistical significance compared to TARDIS but not with Membrain-Seg. Overall, this further demonstrates ETSAM’s ability to consistently outperform TARDIS and ETSAM across Dice, IoU, and Precision, while maintaining high average recall scores.

### 2.2 Comparative Case Study

We conducted a comparative study on ETSAM, Membrain-Seg, and TARDIS-generated membrane segmentation of three cryo-electron tomograms from the test dataset. In Figure 2, for each tomogram, we visually compare the 2D tomogram slice, 3D ground truth membrane annotation (green), alongside predictions from ETSAM (blue), Membrain-Seg (red), and TARDIS (pink). Several key observations emerge from this visualization.

**Fig. 2.**
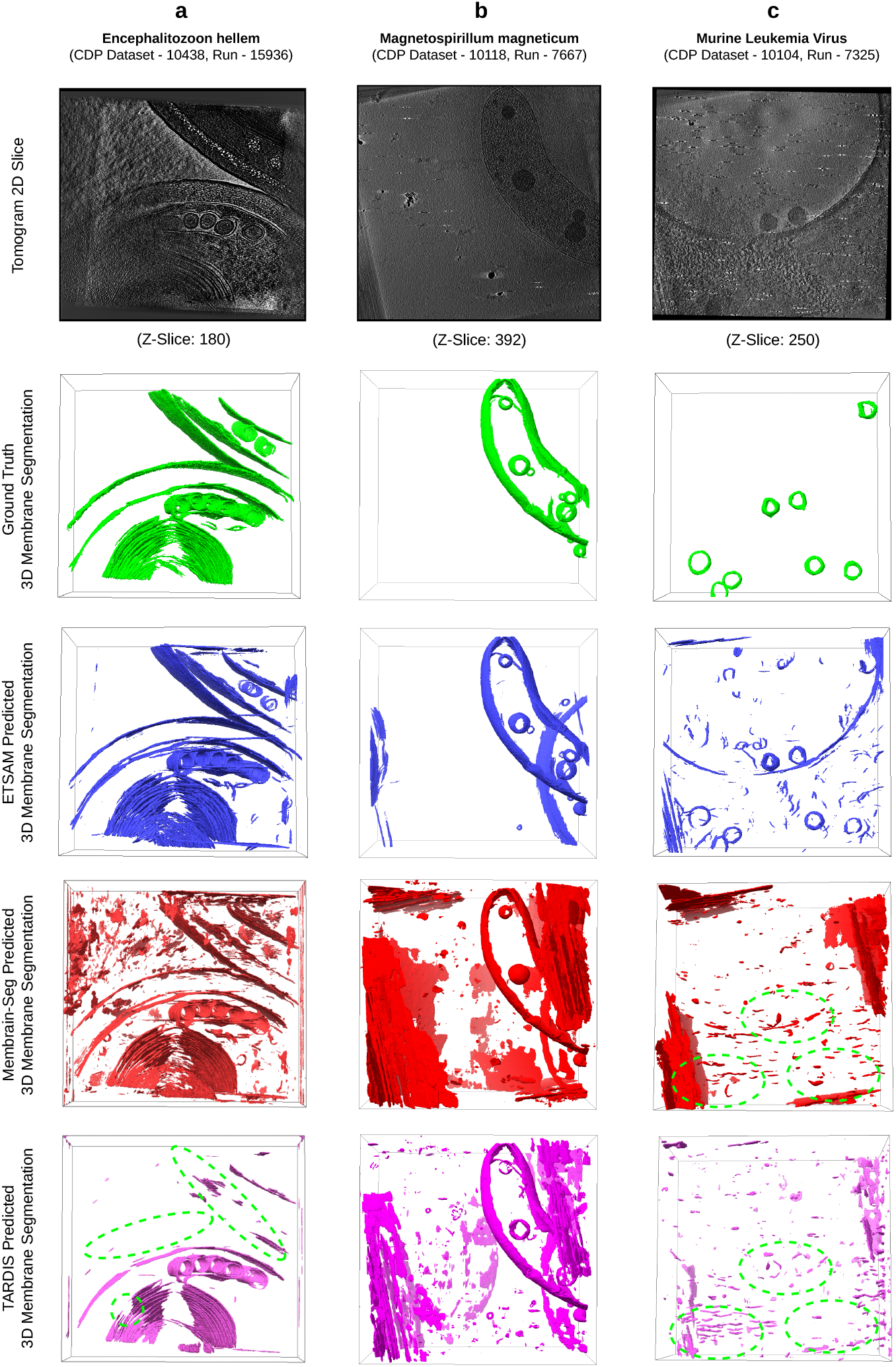
Visual comparison of the ground-truth cell membrane segmentation (green) against the ETSAM (blue), Membrain-Seg (red), and TARDIS (pink) predicted membrane segmentations on three tomograms: **(a)** Encephalitozoon hellem microsporidian parasite cell (CDP Dataset - 10438, Run - 15936), **(b)** Magnetospirillum magneticum bacteria cell (CDP Dataset - 10118, Run - 7667) and **(c)** Murine Leukemia Virus (CDP Dataset - 10104, Run - 7325). Dotted green regions indicate missing ground-truth membranes.

In Figure 2a, we compare the predicted membranes on a tomogram of the Encephal-itozoon hellem microsporidian parasite cell (CDP Dataset - 10438, Run - 15936). ETSAM and Membrain-Seg predict most of the membranes from the ground truth accurately, but TARDIS fails to predict some membranes (green dotted regions). Although Membrain-Seg accurately predicted most ground-truth membranes, it generated a lot of false positive noise. From Supplementary Table 1, we can also statistically observe TARDIS’s low recall (0.4034) scores due to missing membranes and Membrain-Seg’s low precision (0.4347) scores due to false positives. ETSAM’s lower noise (precision: 0.6293) and more complete membrane prediction (recall: 0.7119) help it achieve the best Dice (0.6680) and IoU (0.5015) scores.

In Figure 2b, we compare the predicted membranes on a tomogram of the Magnetospirillum magneticum bacteria cell (CDP Dataset -10118, Run - 7667). Compared with ground-truth membrane annotations (green), all methods accurately predict most membranes, but Membrain-Seg and TARDIS’ (pink) predictions are considerably noisier than those of ETSAM. From Supplementary Table 1, we can note that this leads to a very low precision for Membrain-Seg (0.0827) and TARDIS (0.2626), while ETSAM achieves a precision of 0.7667. This notably improves ETSAM’s Dice (0.8383) and IoU (0.7216) scores compared to the other methods.

Finally, in Figure 2c, we compare the predicted viral envelope or membrane on a tomogram of the Murine Leukemia Virus (CDP Dataset - 10104, Run - 7325). As the virus is surrounded by a lipid bilayer derived from the host cell membrane and thus resembles a cellular membrane, we have included it in our analysis. It can be observed that TARDIS and Membrain-Seg failed to detect the viral membranes (green dotted regions), whereas ETSAM detected most of them accurately. From Supplementary Table 1, we can note that this leads to a very low recall for Membrain-Seg (0.1135) and TARDIS (0.0246). Although ETSAM’s noisy predictions yield lower precision (0.2643), the higher recall (0.6811) indicates that biologically relevant viral membranes were predicted significantly better.

Our comparative analysis highlights ETSAM’s ability to accurately predict biologically relevant membranes in cryo-ET tomograms with significantly lower noise. ETSAM’s cleaner predictions provide better visual clarity into the membrane regions and can significantly reduce manual post-processing and cleaning time for researchers.

### 2.3 Generalization to cryo-ET Tomograms of New Cellular Organisms

To further ensure that ETSAM can robustly segment cell membranes, we also evaluated it on five additional experimental tomograms of different cellular organisms from the CDP database. The cellular organisms in these five experimental tomograms were not part of the training data and have never been seen by ETSAM. Figure 3 visualizes the ETSAM-predicted cell membrane segmentation of these experimental tomograms, along with the respective Membrain-Seg and TARDIS predictions. It can be observed that ETSAM robustly segments cell membranes across various cellular organisms. ETSAM’s predictions contain significantly less noise while predicting membranes on par with Membrain-Seg and TARDIS. Although we cannot quantitatively evaluate ETSAM’s performance due to the lack of ground-truth annotations for these experimental tomograms, the visual comparison clearly demonstrates that it generalizes effectively to new cellular organisms unseen during training.

**Fig. 3.**
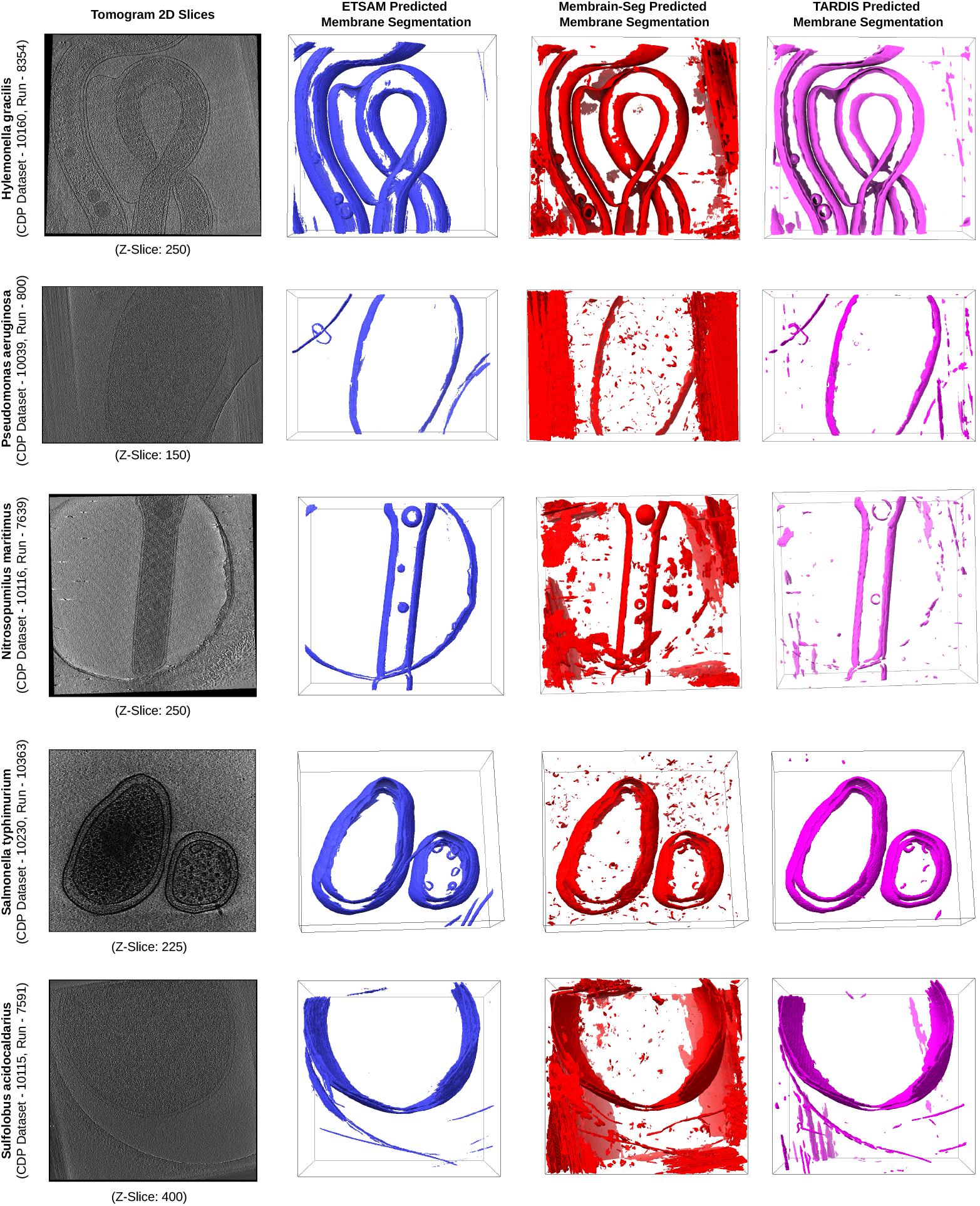
Visual comparison of ETSAM (blue), Membrain-Seg (red), and TARDIS (pink) predicted membrane segmentation across 5 experimental tomograms of different cellular organisms unseen by ETSAM during its training. It demonstrates that ETSAM robustly predicts membranes with less noise than the other two methods.

### 2.4 Post-processing of Membrane Segmentation

During prediction, ETSAM traverses each slice of the tomogram (similar to image frames in a video) to identify membrane boundaries, while its memory encoder and memory attention help track membrane regions across the slices. However, ETSAM may still predict a membrane-like region in a given slice but not observe a corresponding region in subsequent slices, which can introduce thin false-positive noises that increase the difficulty of visualizing and analyzing membrane regions.

To address this drawback, we developed a post-processing technique that reduces visual artifacts in the segmentation results. This is accomplished by identifying 3D blobs in the predicted segmentation mask and removing those that do not extend beyond a specific number (i.e., 10 in this experiment) of consecutive slices. Because the size of noise blobs may vary across tomograms of different sizes, we provide users with an option to adjust the number of slices used to filter the blobs.

We have applied the post-processing technique to all 10 experimental tomograms in our test set. The per-tomogram post-processed ETSAM prediction scores are tabulated in Supplementary Table 2. Figure 4 compares the average scores before and after post-processing using a bar plot. We can observe that post-processing slightly increases the Dice (+0.25%), IoU (+0.31%), and precision (0.51%) scores, with only a marginal decrease in recall (-0.16%). While quantitative improvements are negligible, post-processing can significantly enhance the visual clarity of membrane segmentation by reducing noise, making it easier for users to analyze the results.

**Fig. 4.**
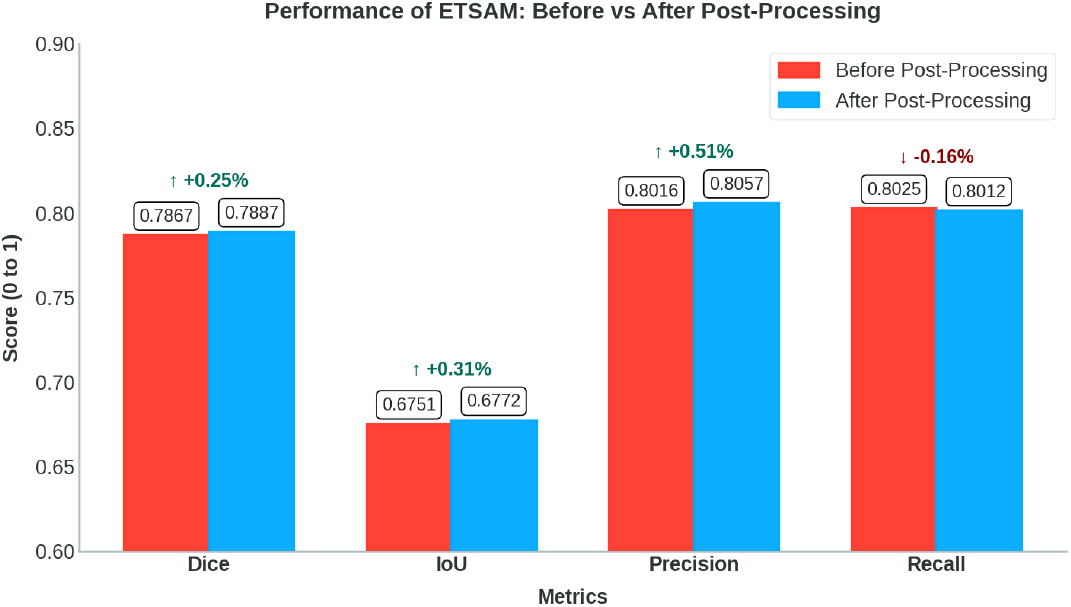
Bar plot that compares the average Dice, IoU, Precision, and Recall scores before and after post-processing ETSAM predicted membranes on the 10 experimental test set tomograms.

Figure 5 visualizes the tomogram of Schizosaccharomyces pombe 972h-strain fungus cell (CDP Dataset - 10000, Run - 247) and the ETSAM predicted segmentation before and after post-processing. It is observed that the post-processing technique removes most noisy regions while preserving the membranes predicted. In this particular tomogram, the post-processing of the ETSAM membrane segmentation improves the Dice score from 0.9595 to 0.9606, the IoU score from 0.9222 to 0.9242, and the precision from 0.9623 to 0.9646 with a negligible reduction in the recall score from 0.9568 to 0.9567. This demonstrates the effectiveness of the post-processing technique in removing noise while preserving biologically relevant membrane regions. Furthermore, Supplementary Fig. 1 presents two additional examples illustrating notable visual improvements achieved through post-processing.

**Fig. 5.**
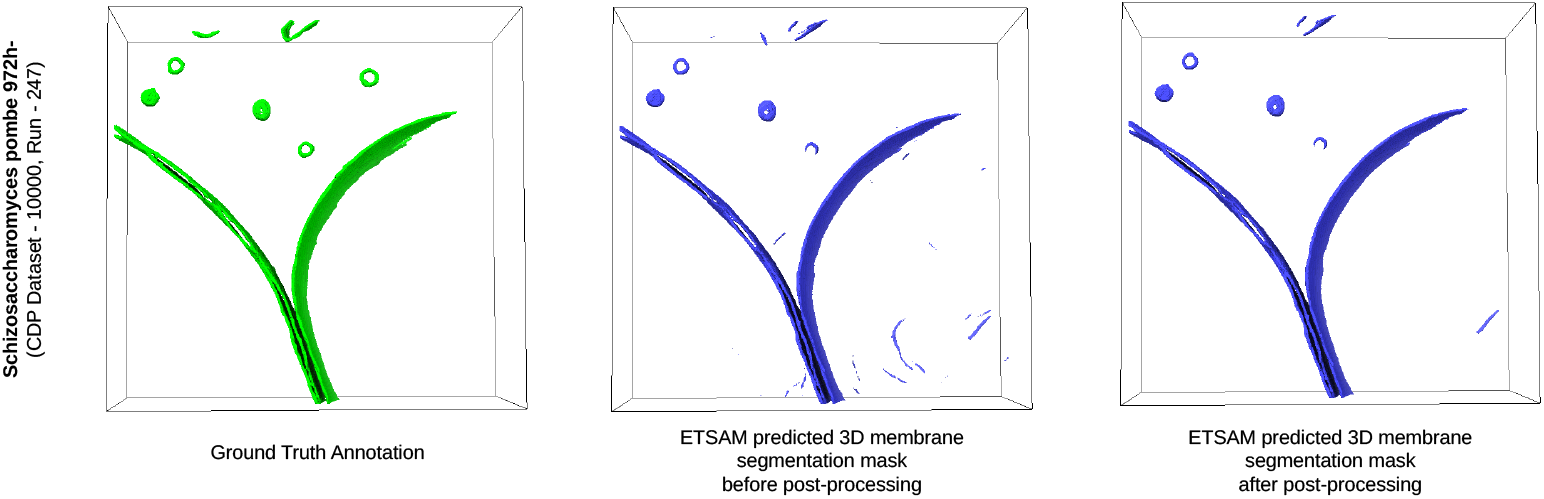
Visualization of the cryo-ET tomogram slice of Schizosaccharomyces pombe 972h- strain fungus cell (CDP Dataset - 10000, Run - 247) along with ETSAM predicted 3D membrane segmentation before and after post-processing.

While post-processing can improve the visual clarity of ETSAM membrane segmentation, caution is warranted when processing tomograms with thin or small membrane regions, as they may be removed. We recommend visually comparing unprocessed and post-processed segmentation results to ensure biologically relevant membrane regions are not removed.

### 2.5 Runtime and Compute Benchmark

A single cryo-EM grid may contain hundreds of square patches, each with dozens to hundreds of holes to house the specimens. Modern automated, multi-shot acquisition techniques enable high-throughput tilt-series acquisition, where processing and analysis become the bottleneck. This necessitates rigorous runtime and compute benchmarking of tomogram segmentation techniques to ensure scalable data processing and analysis. We compared the runtime, CPU memory, and GPU memory usage of ETSAM, Membrain-Seg, and TARDIS on the 10 experimental tomograms in the test set. The per-tomogram results are displayed in Figure 1d and tabulated in Supplementary Table 3. ETSAM achieves the fastest average runtime (70.81 ± 28.67 seconds per tomogram), followed by Membrain-Seg (90.27 ± 43.05 seconds per tomogram) and TARDIS (383.54 ± 509.13 seconds per tomogram). In terms of average CPU memory, TARDIS consumes the least (9661.1 ± 6577.0 MB per tomogram), followed by ETSAM (16453.6 ± 6636.7 MB) and Membrain-Seg (17802.7 ± 7791.2 MB). CPU memory usage appears to vary with the size of the tomogram. Crucially, the GPU memory consumption of ETSAM (1354.6 ± 24.2 MB) is 3.88x lower than Membrain-Seg (5252.0 ± 713.4 MB) and 5.84x lower than TARDIS (7906.0 ± 0.0 MB). Since ETSAM processes a tomogram slice by slice, it uses GPU memory more efficiently than other methods. Overall, the lower GPU memory footprint and faster runtime make ETSAM the ideal choice for high-throughput workloads. It also enables running it on laptops/desktops with smaller GPUs. Benchmarks were run on an Intel® Core™ i7-14700K CPU, 32 GB DDR5 RAM, and an Nvidia RTX 4070 GPU with 12 GB of VRAM.

### 2.6 Limitations

ETSAM’s high sensitivity effectively predicts most membrane regions, but it can also misclassify some membrane-like artifacts as membranes, introducing noise into the predictions. These membrane-like artifacts can occur during cryo-ET sample preparation, imaging, or tomogram reconstruction. In the case of cryo-electron tomography of Schizosaccharomyces pombe 972h-strain fungus cell (CDP Dataset - 10001, Run - 251), as shown in Supplementary Fig. 2a, ETSAM, Membrain-Seg, and TARDIS predict a membrane-like ice-contamination artifact (false-positive) that is not an actual cell membrane. Supplementary Fig. 2b shows another example in which ETSAM, Membrain-Seg, and TARDIS predict border noises likely caused by geometric artifacts [46] as membranes to varying extents in the tomogram of a Halobacterium salinarum bacteria cell (CDP Dataset - 10099, Run - 7042). In Supplementary Fig. 2c, the visualization of the orange dotted region in ETSAM prediction shows that it falsely predicts cryo-EM grid carbon film holes as membranes in the tomogram of Nitrosopumilus maritimus archaeon cell (CDP Dataset - 10116, Run - 7639). TARDIS and Membrain-Seg appear to handle it better than ETSAM in this case. These types of false-positive membrane predictions require manual inspection and cleaning. In the future, larger training datasets with high-quality ground-truth annotations and more advanced AI models can help mitigate these shortcomings. Technological and scientific advancements in cryo-ET sample preparation, imaging, and tomogram reconstruction can also significantly reduce the artifacts introduced in tomograms.

## 3 Conclusion

Accurate identification of cell membranes in cryo-ET tomograms is essential for understanding cellular architecture and function, yet remains challenging due to low signal-to-noise ratios, reconstruction artifacts, and the substantial effort required for manual annotation in large 3D volumes. In this study, we introduced ETSAM, a two-stage SAM2-based model that delivers fully automated, state-of-the-art membrane segmentation in cryo-ET tomograms. ETSAM achieved the best overall performance across all evaluation metrics and offered a superior balance between precision and recall compared with existing deep learning approaches, as evidenced by its substantially higher AUPRC score on the experimental test tomograms. Its strong recall facilitates the detection of biologically relevant membrane regions, while its high precision reduces noise and false positives, thereby reducing the time researchers spend on manual cleaning and correction. In addition, its fast runtime and low GPU memory footprint enable high-throughput processing and ease of access. Together, these capabilities position ETSAM as a robust, practical tool for advancing structural and cellular studies using cryo-ET.

## 4 Methods

### 4.1 Data Collection

The experimental cryo-ET tomograms and corresponding cell membrane annotations used in this work were collected from the CDP database [44]. To create the training dataset, we manually reviewed the experimental cryo-ET tomograms in the CDP database and selected 83 tomograms from 25 CDP datasets with properly annotated cell membranes. We also curated a test dataset comprising 10 experimental tomograms from 8 CDP datasets to evaluate ETSAM’s performance. Additionally, we collected 5 experimental tomograms of different cellular organisms from the CDP database. These tomograms were used to visually validate that ETSAM could robustly handle tomograms of new cellular organisms unseen during training.

In addition to the experimental tomograms curated from the CDP database, we used PolNet to generate 28 simulated tomograms. These simulated tomograms were included in the training dataset. In total, the curated training and test datasets contain 111 and 10 tomograms, respectively.

### 4.2 Dataset Preparation

To enhance the quality of the training dataset, we manually cleaned the cell membrane annotations of the experimental tomograms to remove noise and false-positive membrane segmentation, as illustrated in Supplementary Fig. 3. Moreover, since the density distributions of the experimental tomograms vary, we normalize them to the range 0.0 to 1.0 using min-max normalization.

As SAM2 [43] is designed to take 2D RGB image frames from videos as input, ETSAM also treats a 3D tomogram as a video and slices it along the Z-axis to produce image-like frames. Since an image is composed of RGB channels, ETSAM repeats each 2D slice of tomogram data across three channels to form the RGB channels in an image. Similarly, the 3D cell membrane annotation masks are also converted into 2D slices. Finally, the 2D image-like tomogram and corresponding membrane annotation slices are stored in input-label pairs to train the ETSAM model. Although we convert 3D tomograms into 2D image-like slices, it is worth noting that the tomogram data is not limited to 0-255 integer values like in traditional RGB images. Instead, they are stored as floating-point values (0.0 to 1.0) in NumPy (.npy) format, thereby preserving the precision of the tomogram data.

For our test dataset, we manually created ground-truth annotations using a hybrid approach. A combination of existing annotations from the CDP database and annotations generated by AI models such as TARDIS, Membrain-Seg, and ETSAM was used as initial and reference annotations. To generate the ground-truth annotation, the initial annotations were extensively cleaned to remove noise, manually corrected slice by slice to ensure accuracy, and compared with the reference annotations to prevent missing membranes. In Supplementary Fig. 4, we illustrate two examples of the initial annotations that were manually cleaned, corrected, and combined to form the ground truth annotations.

### 4.3 ETSAM Membrane Segmentation Pipeline

The ETSAM pipeline consists of two SAM2-based [43] blocks to perform two-stage membrane segmentation, as shown in Figure 6a. While the original SAM2 model was designed to accept RGB images (.jpg/.png) with an integer value range of 0 to 255, ETSAM customizes it to load tomogram slices with floating-point values in the range [0.0, 1.0]. The input tomograms are normalized to the range by using min-max normalization. The normalized tomogram slices are first fed into the ETSAM Stage 1 block, which generates the initial membrane prediction. The first-stage segmentation is in logit form, with values ranging from negative to positive, where higher values indicate a greater likelihood of membrane presence. The predicted logits are then clipped to the range -5 to +5 and normalized via min-max scaling. These normalized first-stage predicted logits are then concatenated with the normalized tomogram slices and fed as input to the ETSAM Stage 2 block, which generates the final membrane prediction.

**Fig. 6.**
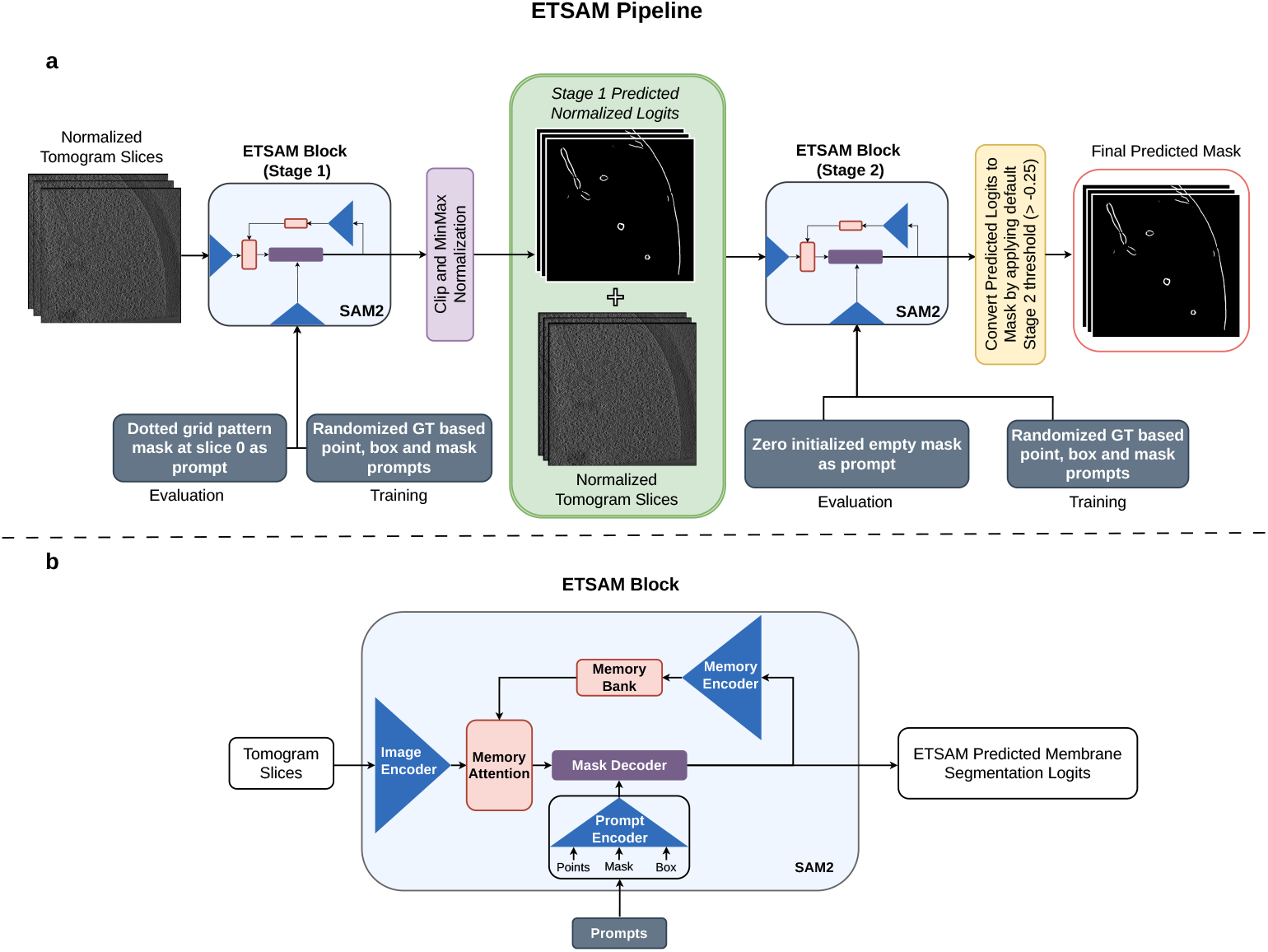
Overview of ETSAM. **(a)** The pipeline of the two-stage ETSAM. **(b)** The network architec- ture of the ETSAM block is used in both stage 1 and stage 2.

As shown in Figure 6b, each ETSAM stage block features a SAM2 comprising an image encoder, a prompt encoder, a mask decoder, and specialized memory components, such as a memory encoder and memory attention. The image encoder serves as the foundational component for feature extraction, using a Hiera (Hierarchical transformer-based architecture) [47] that processes input frames at multiple resolutions to capture both local details and global context. Unlike traditional Vision Transformers (ViT), Hiera employs a multi-stage design with quadtree-like down-sampling, allowing for efficient hierarchical feature maps that reduce computational overhead while preserving spatial hierarchies. For video inputs, the encoder processes each frame independently, generating per-frame embeddings that form the basis for subsequent prompting and decoding.

The prompt encoder encodes user-provided prompts into the model’s latent space. It handles various prompt types — points, bounding boxes, and masks — by embedding them into high-dimensional vectors. In the prompt encoder, points and box prompts are encoded using positional embeddings, and mask prompts are passed through convolutional layers and positional encoding to align with the image encoder’s features. The memory encoder is a lightweight convolutional network that compresses mask predictions from previous frames into compact representations using fixed-dimensional tokens that capture essential spatial and semantic information. These tokens are then stored in the memory bank. A memory bank is a bounded buffer that stores representations of a fixed number of recent frames, using a first-in-first-out (FIFO) strategy to manage the memory footprint. Finally, the mask decoder is a lightweight bidirectional transformer with cross-attention and self-attention layers that integrate the encoded prompts with the image and memory bank features to predict the segmentation mask.

Both the ETSAM’s Stage 1 and 2 were initialized with the SAM2.1 base model checkpoint as the initial weights and fine-tuned on the training dataset for 602 and 545 epochs, respectively. During training, ETSAM uses the default random ground-truth-based prompting configuration from SAM2. It generates points, box, and mask prompts sampled from the ground-truth annotations with probabilities of 0.5, 0.25, and 0.25, respectively. ETSAM was trained on NVIDIA A100 with a batch size of 1. Each batch corresponds to 22 consecutive 2D slices from a single tomogram. Overall, both stages of ETSAM use the same default model parameters and training regime as the original SAM2.

During inference, as shown in Figure 6a, the ETSAM stage 1 uses a fixed grid of points mask (dotted grid pattern) at the first slice as the input prompt, and an empty mask initialized with zeros is used as input prompt for ETSAM Stage 2. The prompting process for both stages is fully automated and requires no user input. Finally, the logits predicted by the second-stage ETSAM block are converted into a binary segmentation mask by applying a default logit threshold of −0.25. Users can adjust this threshold to balance the precision and recall of the final prediction.

### 4.4 Ablation Studies of Prompting Techniques and One/Two-Stage Segmentation

SAM2 [43] is a foundational AI image/video segmentation model that allows users to provide manual input prompts to segment and track regions of interest. However, we want ETSAM to operate in a fully automated manner, similar to other competing methods that require no manual user input during prediction. To address this problem, in our first ablation study, we tested three prompting techniques that automatically generate custom generic input prompts for ETSAM Stage 1.

The first prompting technique initializes an empty mask with zeroes in the first slice as the input prompt. The second prompting technique uses a mask-based prompt initialized with a grid of points (dotted grid pattern) spaced by 50 pixels in the XY plane of the first slice as the input prompt. The third prompting technique is similar to the second except that the dotted grid pattern is repeated in the first and every 50^*th*^ slice of the input prompt mask. For each pixel in a given tomogram slice, the final layer of ETSAM’s mask decoder predicts a logit value. The predicted logits can range from negative to positive values, with higher values indicating a greater likelihood that a cell membrane is present at that pixel. We used a default logit threshold of 0.0 for converting ETSAM Stage 1 predicted logits into binary segmentation masks, where any voxel with a logit value greater than the threshold is segmented as a cell membrane. These binary segmentation masks were then used for evaluation.

ETSAM Stage 1 was evaluated using the three prompting techniques on the 10 experimental tomograms in the test dataset, and the results are tabulated in Supplementary Table 4. The first prompting technique of initializing an empty mask prompt performs poorly and fails to generate proper segmentation masks in some cases. The second prompting technique of initializing the grid of points at only the first slice achieves significantly better results than the other two approaches. This approach enables the first stage of ETSAM to initially concentrate equally on all regions, and as membrane regions are identified, it tracks them continuously using memory features without interruption. This is unlike the third approach, which is interrupted by a grid prompt every 50^*th*^ slice and leads to poorer performance. Therefore, the second automated prompting technique, which achieved the best results, is selected as the default for Stage 1.

In the two-stage ETSAM pipeline, as shown in Figure 6, the ETSAM Stage 1 predicted logits are clipped between -5 and 5, then min-max normalized to the 0 to 1 range, and added back to the original tomogram as input to ETSAM Stage 2. The ETSAM Stage 2 predicted logits are finally converted into a binary membrane segmentation mask using a logit threshold. In the second ablation study, we conducted a comprehensive evaluation of various logit thresholds and prompting techniques for ETSAM Stage 2 on the experimental tomograms in our test set.

In addition to testing the same three prompting techniques used in Stage 1, we developed a fourth prompting technique specifically for Stage 2. The fourth prompting technique uses the Stage 1 predicted segmentation mask (generated with 0.0 threshold) of the first and every 50^*th*^ slice as the input prompt for Stage 2. For each of the four prompting techniques, we tested nine logit thresholds: [-2, -1, -0.5, -0.25, 0, 0.25, 0.5, 1, 2]. The per-tomogram results are tabulated in Supplementary Tables 5, 6, 7, and 8. It can be observed that using a zero-initialized empty mask with a logit threshold of -0.25 performs best on average and is therefore selected as the default for ETSAM Stage 2. It also shows that no single logit threshold performs best across all tomograms. Therefore, we also provide users with the option to store the prediction as logits, which can be thresholded after careful visualization and analysis. Thus, allowing users to achieve the preferred precision-recall trade-off on a per-tomogram basis.

In our final ablation study, we compared ETSAM with single- and two-stage architectures on the test dataset; per-tomogram results are reported in Supplementary Tables 9 and 1, respectively. We can observe that the two-stage ETSAM outperforms the single-stage ETSAM across all metrics. The improvements in both precision and recall in two-stage ETSAM indicate that it can detect more membranes while reducing false positives. The three examples illustrated in Supplementary Fig. 5 show that the two-stage ETSAM also substantially improves the visual clarity of the predicted membrane segmentation with reduced random and border noises. Although adjusting the logit thresholds can improve precision, it comes at the expense of lower recall. In contrast, ETSAM Stage 2 can improve Stage 1 predictions in terms of both recall and precision while also yielding notably less noisy results. This highlights the benefit of the two-stage ETSAM pipeline introduced in this study.

### 4.5 Evaluation Metrics

In the context of binary segmentation of cell membranes in a tomogram, True Positives (TP) are the correctly predicted positive pixels/membrane regions, False Positives (FP) are the regions incorrectly predicted as membranes (noise/over-segmentation), False Negatives (FN) are the missed membrane regions and True Negatives (TN) are the background (non-membrane) regions predicted correctly.

**Precision** measures the accuracy of a model’s positive prediction and is computed as the ratio of true positives to the total true and false positives. It ranges from 0 to 1, with 1 representing the highest precision, characterized by no false positive predictions, and 0 representing the lowest precision, where none of the positive regions are predicted correctly. It is calculated by the following equation:

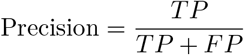

**Recall** measures the completeness of the model’s positive predictions and is computed as the ratio of true positives to the sum of true positives and false negatives. It ranges from 0 to 1, with 1 indicating the highest recall (all positive regions are predicted correctly) and 0 indicating the lowest recall (none of the positive regions are predicted). It is calculated as follows:

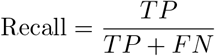

**Dice Score**, also known as **F1 score**, is the harmonic mean of precision and recall, providing a single metric that balances both the accuracy of the prediction and its completeness. It ranges from 0 (no overlap) to 1 (perfect overlap) and penalizes extreme values, i.e., if either precision or recall is low, the Dice score will suffer significantly. Given two sets, *A* (prediction) and *B* (ground truth), it is calculated as follows:

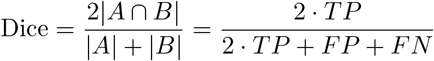

**Intersection over Union (IoU)** is the ratio of the area of overlap between the predicted segmentation and the ground truth to the area of their union, measuring the extent to which the two distinct sets share the same space. It ranges from 0 (no overlap) to 1 (perfect overlap), similar to the Dice score, but is slightly more stringent and effectively penalizes both false positives and false negatives. Given two sets, *A* (prediction) and *B* (ground truth), it is calculated as follows:

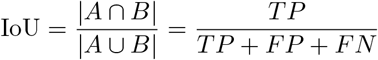

**Area Under Precision-Recall Curve (AUPRC)** measures the area of the Precision-Recall (PR) curve. The PR curve plots Precision (*y*-axis) against Recall (*x*-axis) across various decision thresholds between 0 and 1. If the model-predicted probability at a given tomogram voxel exceeds the decision threshold, it is classified as a positive prediction; otherwise, it is classified as a negative prediction. AUPRC ranges from 0 to 1. A higher AUPRC score indicates that a model achieves overall higher recall and precision across different decision thresholds.

While TARDIS provides an option to output per-voxel probability scores, Membrain-Seg and ETSAM only provide the logits from the model’s last layer (before thresholding). Membrain-Seg predicted logits are converted into probability scores by applying a sigmoid function. In the case of ETSAM, to prevent extreme negative values (e.g., -1024) from skewing the probability distribution, we clip the negative values to the maximum positive value in the predicted logits. We then use a sigmoid function to convert the clipped logits into probability scores.

#### Paired Student’s T-test

Student’s t-test is a statistical test used to determine whether the difference between the scores or results of two groups is statistically significant.

To compare three membrane segmentation models in this work, we conducted 3 separate paired Student’s t-tests as follows: Model A vs Model B, Model B vs Model C, and Model C vs Model A. However, at a significance level of 0.05 (i.e., confidence level 95%), each paired t-test incurs a 5% chance of error, leading to a higher error rate than the expected significance level of 0.05. To correct this, we can apply the Bonferroni correction to the significance level by dividing it by the number of paired comparisons. Thus, in the case of a 3 model comparison, to achieve a significance level of 0.05, the Bonferroni corrected significance level must be calculated as follows: *α* = 0.05/3 = 0.0167. If any given pair’s *p* − *value* < 0.0167, then they are statistically significant, and if *p* − *value* > 0.0167, there is no statistical significance.

**Shapiro-Wilk test** is a statistical test used to determine if a sample of continuous data comes from a normally distributed population. It is used to check the normality assumption before performing a t-test. It works by comparing the sample data to a normal distribution, yielding a p-value, where:

If (*p* > 0.05) - Data is likely normal; proceed with the t-test.

If (*p* < 0.05) - Data is not normal; indicates a significant departure from normality.

Not suitable for the t-test.

All the above evaluation metrics and methods were used to provide a comprehensive analysis of ETSAM’s ability to effectively segment cell membranes in cryo-ET tomograms.

## Supplementary information

We have included Supplementary Tables and Supplementary Figures.

## Acknowledgements

This work is supported by a National Institutes of Health (NIH) grant (grant number: R01GM146340).

## Declarations

## Funding

This work is supported by a National Institutes of Health (NIH) grant (grant number: R01GM146340).

## Conflict of interest/Competing interests

The authors declare no competing interests.

## Data availability

The annotations and simulated tomograms used in training and test datasets are available in Harvard Dataverse. The experimental tomograms can be fetched from the CryoET Data Portal. We have also provided helper scripts to download the entire dataset and preprocess it in our GitHub repository.

## Code availability

The source code of ETSAM is freely available in GitHub. ETSAM model weights are available in Zenodo.

## Author contribution

J.C. conceived the project; J.S. collected the data and carried out the experiments under the guidance of J.C.; J.S. and J.C. analyzed the results; J.S. and J.C. wrote the manuscript.

## Notes

https://github.com/jianlin-cheng/ETSAM/

https://zenodo.org/records/17571925

https://dataverse.harvard.edu/dataset.xhtml?persistentId=doi:10.7910/DVN/K4JKCW

### Competing Interest Statement

The authors have declared no competing interest.

### Summary of Updates

Correct the reference of PolNet tool.

