## Supplementary Figures for "ETSAM: Effectively Segmenting Cell Membranes in cryo-Electron Tomograms"

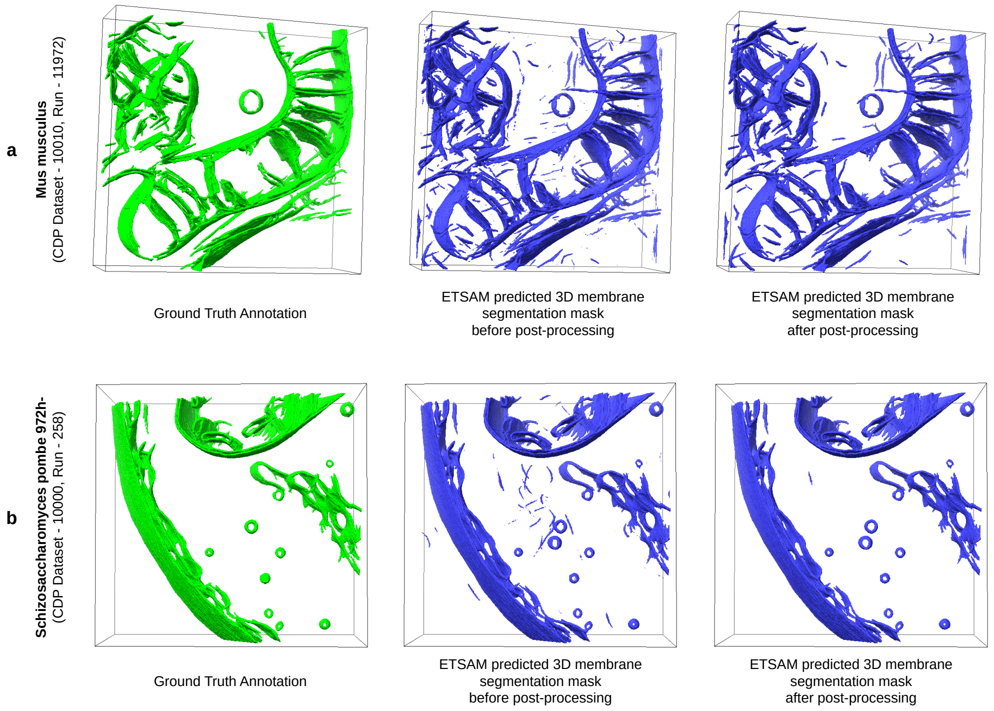


**Fig. 1** Visual improvements due to post-processing ETSAM Membrane Segmentation predicted on cryo-electron tomograms of **(a)** Mus musculus's embryonic fibroblast cell (CDP Dataset - 10010, Run - 11972) and **(b)** Schizosaccharomyces pombe 972h- strain fungus cell (CDP Dataset - 10000, Run - 258).

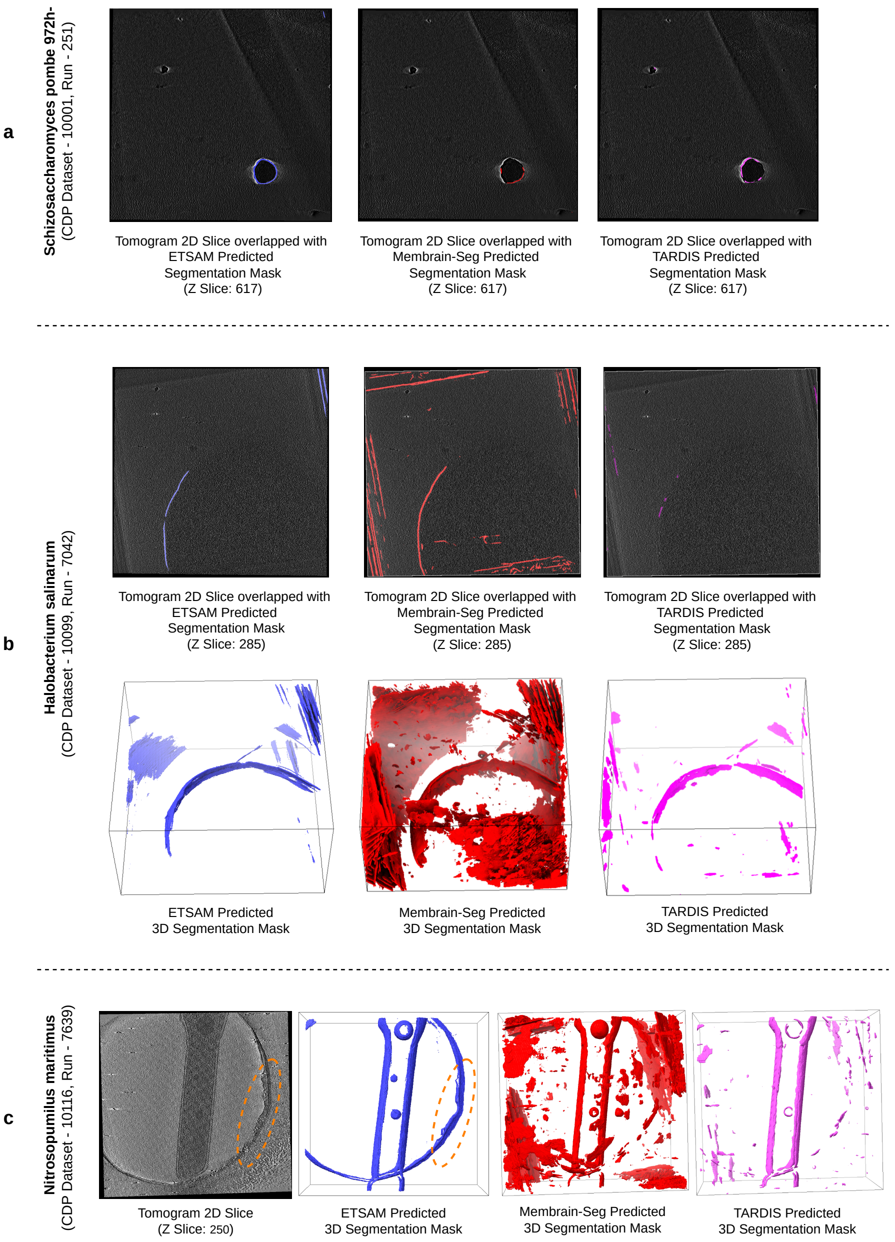


**Fig. 2** Examples showing few limitations of deep learning methods in handling noise artifacts. **(a)** Shows membrane-like artifacts being predicted as a membrane region by ETSAM (blue), Membrain-Seg (red), and TARDIS (pink) in the bottom right of the tomogram slice of Schizosaccharomyces pombe 972h- strain fungus cell (CDP Dataset - 10001, Run - 251). **(b)** Compares border noise likely caused by geometric artifacts being predicted as membrane regions to various extents by ETSAM (blue), Membrain-Seg (red), and TARDIS (pink) in the tomogram of the Halobacterium salinarum bacteria cell (CDP Dataset - 10099, Run - 7042). **(c)** Shows cryo-EM grid carbon film hole (orange dotted region) in the tomogram of the Nitrosopumilus maritimus archaeon cell (CDP Dataset - 10116, Run - 7639) being detected as membrane by ETSAM (blue) to a large extent, while Membrain-Seg (red) and TARDIS (pink) are only partially affected by it.

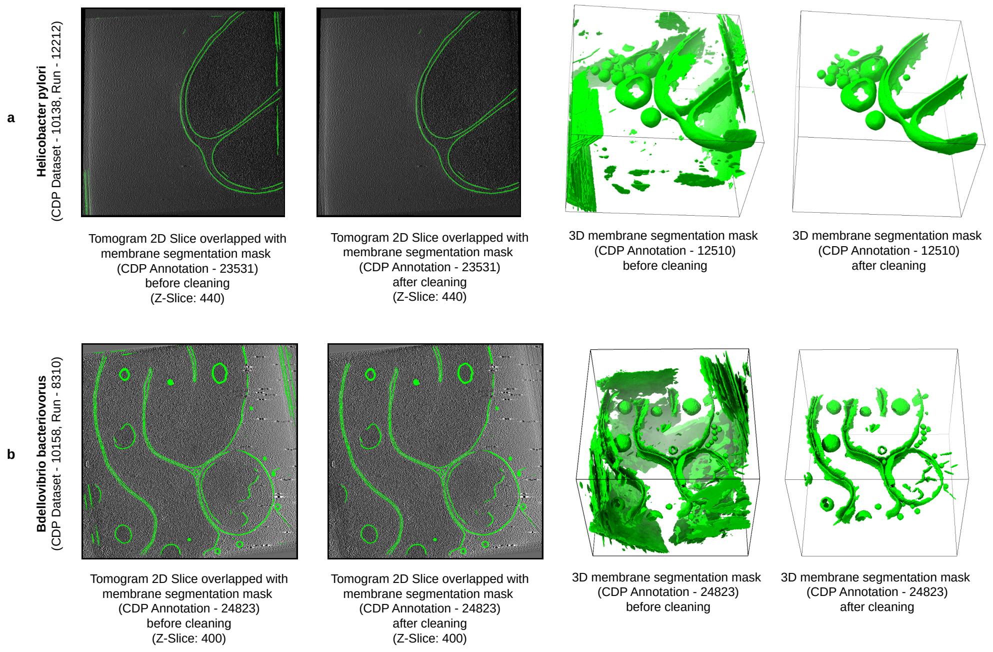


**Fig. 3** Visualization of the improvements due to manually processing membrane annotations of two experimental tomograms from our training dataset. **(a)** Compares the membrane segmentation of the Helicobacter pylori bacteria cell (CDP Dataset - 10138, Run - 12212) before and after cleaning random and geometric border artifacts. **(b)** Compares the membrane segmentation of the Bdellovibrio bacteriovorus bacteria cell (CDP Dataset - 10158, Run - 8310) before and after cleaning the geometric border artifacts.

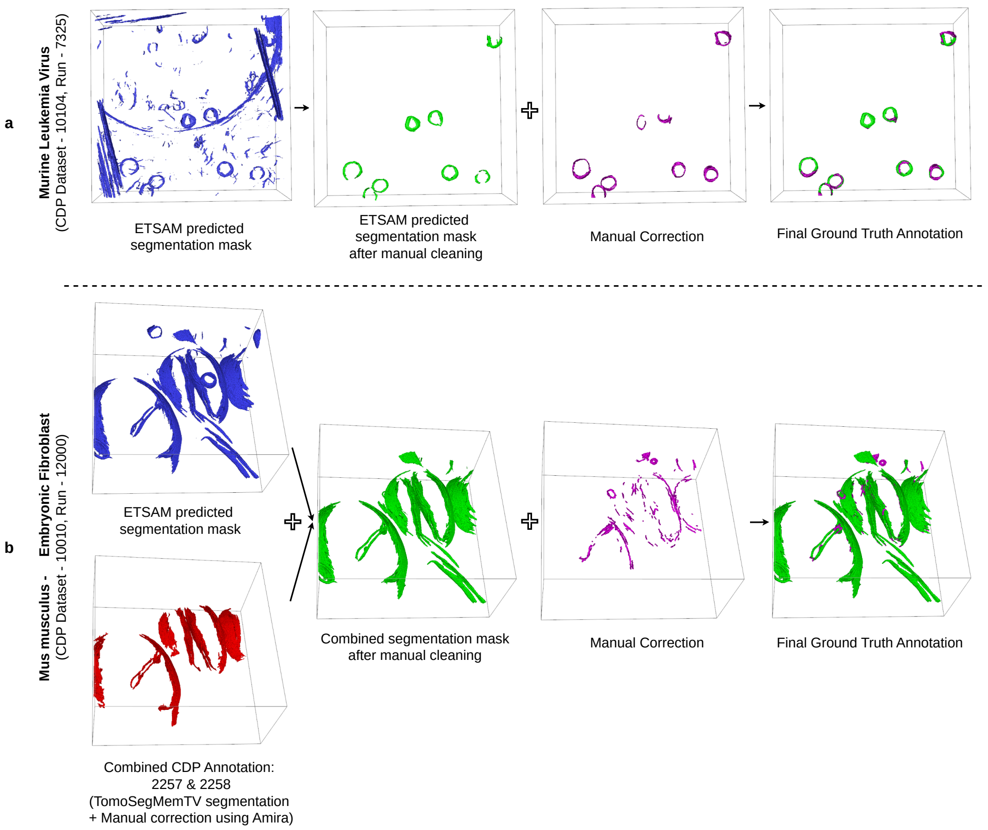


**Fig. 4** Visualization of the hybrid approach used for creating ground-truth annotations of two experimental tomograms from our test dataset. **(a)** Shows the initial ETSAM predicted membrane annotation being manually cleaned and corrected to form the ground-truth annotation for a tomogram of Murine Leukemia Virus (CDP Dataset - 10104, Run - 7325). **(b)** Shows the initial ETSAM predicted membrane annotation for a tomogram of Mus musculus's embryonic fibroblast cell (CDP Dataset - 10010, Run - 12000) being combined with existing CDP Annotation (2257 & 2258) and manually cleaned and corrected to form the ground-truth annotation.

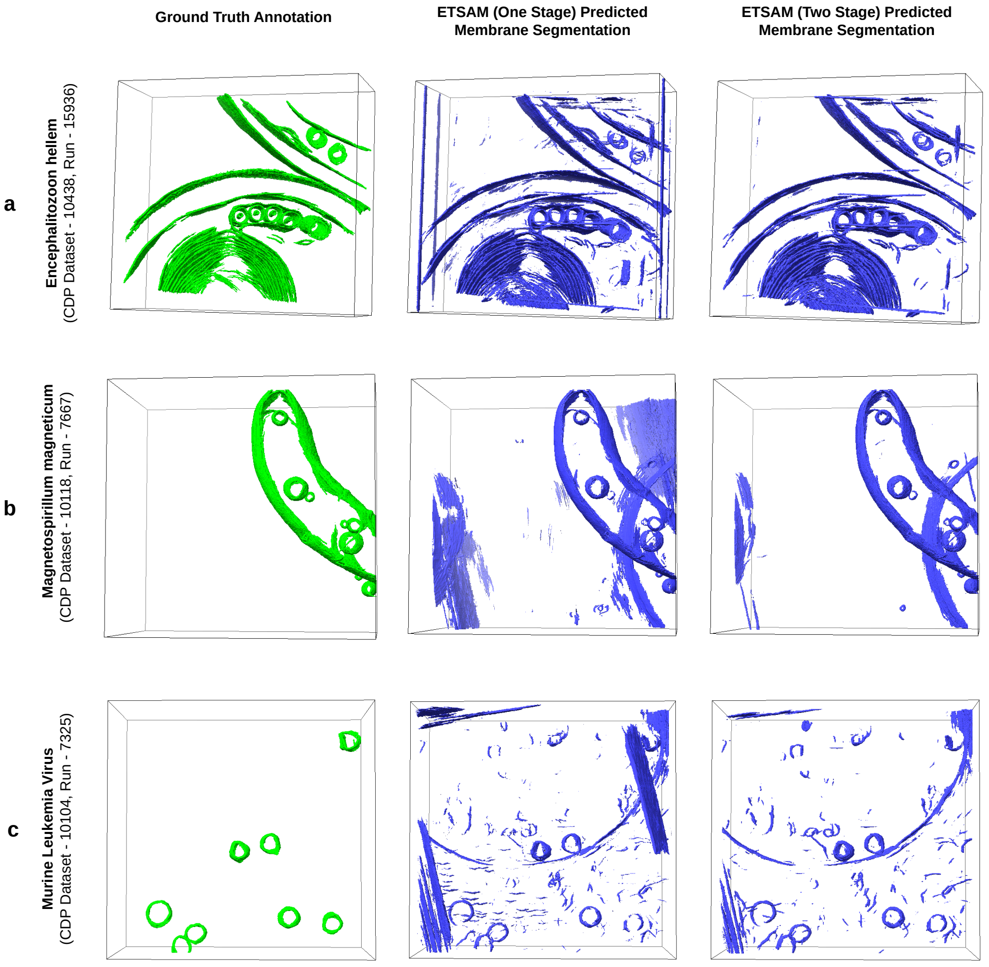


**Fig. 5** Comparison between Ground Truth Annotation, ETSAM (one stage), and ETSAM (two stage) predicted membrane segmentation on experimental tomograms of **(a)** Encephalitozoon hellem microsporidian parasite cell (CDP Dataset - 10438, Run - 15936), **(b)** Magnetospirillum magneticum bacteria cell (CDP Dataset - 10118, Run - 7667), and **(c)** Murine Leukemia Virus (CDP Dataset - 10104, Run - 7325).
