## Supplementary Tables for "ETSAM: Effectively Segmenting Cell Membranes in cryo-Electron Tomograms"

**Supplementary Table 1:** Per-Tomogram Performance of two-stage ETSAM on experimental tomograms in the test set

| CDP IDs | | ETSAM | | | | TARDIS | | | | Membrain-Seg | | | |
|---|---|---|---|---|---|---|---|---|---|---|---|---|---|
| **Dataset** | **Run** | **Dice/F1** | **IoU** | **Precision** | **Recall** | **Dice/F1** | **IoU** | **Precision** | **Recall** | **Dice/F1** | **IoU** | **Precision** | **Recall** |
| 10000 | 245 | 0.7261 | 0.5700 | 0.8780 | 0.6191 | 0.5706 | 0.3992 | 0.9418 | 0.4093 | 0.7512 | 0.6015 | 0.7360 | 0.7670 |
| 10000 | 247 | 0.9595 | 0.9222 | 0.9623 | 0.9568 | 0.8139 | 0.6862 | 0.8291 | 0.7992 | 0.7875 | 0.6494 | 0.6837 | 0.9283 |
| 10001 | 258 | 0.9519 | 0.9082 | 0.9645 | 0.9396 | 0.7496 | 0.5995 | 0.9275 | 0.6289 | 0.8225 | 0.6985 | 0.8344 | 0.8109 |
| 10010 | 11972 | 0.8982 | 0.8152 | 0.8856 | 0.9112 | 0.7049 | 0.5443 | 0.8439 | 0.6052 | 0.7730 | 0.6300 | 0.7783 | 0.7679 |
| 10010 | 12000 | 0.9041 | 0.8250 | 0.9014 | 0.9068 | 0.6343 | 0.4645 | 0.5171 | 0.8203 | 0.6128 | 0.4418 | 0.4749 | 0.8635 |
| 10438 | 15936 | 0.6680 | 0.5015 | 0.6293 | 0.7119 | 0.5069 | 0.3395 | 0.6818 | 0.4034 | 0.5263 | 0.3571 | 0.4347 | 0.6668 |
| 10436 | 15943 | 0.7671 | 0.6222 | 0.8208 | 0.7200 | 0.5028 | 0.3358 | 0.6014 | 0.4320 | 0.5214 | 0.3526 | 0.4194 | 0.6890 |
| 10104 | 7325 | 0.3809 | 0.2352 | 0.2643 | 0.6811 | 0.0162 | 0.0081 | 0.0120 | 0.0246 | 0.0078 | 0.0039 | 0.0040 | 0.1135 |
| 10118 | 7667 | 0.8383 | 0.7216 | 0.7667 | 0.9246 | 0.4056 | 0.2544 | 0.2626 | 0.8901 | 0.1503 | 0.0812 | 0.0827 | 0.8195 |
| 10032 | 676 | 0.7726 | 0.6294 | 0.9430 | 0.6543 | 0.6485 | 0.4798 | 0.5842 | 0.7286 | 0.4459 | 0.2869 | 0.2927 | 0.9357 |
| **AVERAGE** | | 0.7867 | 0.6751 | 0.8016 | 0.8025 | 0.5553 | 0.4111 | 0.6201 | 0.5742 | 0.5399 | 0.4103 | 0.4741 | 0.7362 |

**Supplementary Table 2:** Per-Tomogram Performance of two-stage ETSAM on experimental tomograms in test set after post-processing

| CDP IDs | | ETSAM | | | |
|---|---|---|---|---|---|
| **Dataset** | **Run** | **Dice/F1** | **IoU** | **Precision** | **Recall** |
| 10000 | 245 | 0.7240 | 0.5674 | 0.8788 | 0.6156 |
| 10000 | 247 | 0.9606 | 0.9242 | 0.9646 | 0.9567 |
| 10001 | 258 | 0.9536 | 0.9114 | 0.9684 | 0.9393 |
| 10010 | 11972 | 0.9002 | 0.8185 | 0.8895 | 0.9112 |
| 10010 | 12000 | 0.9049 | 0.8263 | 0.9058 | 0.9041 |
| 10438 | 15936 | 0.6687 | 0.5023 | 0.6308 | 0.7116 |
| 10436 | 15943 | 0.7670 | 0.6220 | 0.8236 | 0.7176 |
| 10104 | 7325 | 0.3951 | 0.2462 | 0.2784 | 0.6803 |
| 10118 | 7667 | 0.8416 | 0.7265 | 0.7725 | 0.9242 |
| 10032 | 676 | 0.7709 | 0.6272 | 0.9444 | 0.6513 |
| **AVERAGE** | | 0.7887 | 0.6772 | 0.8057 | 0.8012 |

**Supplementary Table 3:** Benchmark of runtime, CPU and GPU memory compute usage of all three methods evaluated on experimental tomograms in the test set

| CDP IDs | | ETSAM | | | TARDIS | | | Membrain-Seg | | |
|---|---|---|---|---|---|---|---|---|---|---|
| **Dataset** | **Run** | **Time (seconds)** | **GPU Memory (MB)** | **CPU Memory (MB)** | **Time (seconds)** | **GPU Memory (MB)** | **CPU Memory (MB)** | **Time (seconds)** | **GPU Memory (MB)** | **CPU Memory (MB)** |
| 10000 | 245 | 72.93 | 1350 | 20582 | 451.08 | 7906 | 8131 | 98.95 | 5520 | 20985 |
| 10000 | 247 | 73.36 | 1350 | 21160 | 129.19 | 7906 | 8328 | 104.60 | 5520 | 21169 |
| 10001 | 258 | 68.34 | 1350 | 21130 | 123.01 | 7906 | 7841 | 94.73 | 5520 | 21212 |
| 10010 | 11972 | 52.13 | 1348 | 9853 | 67.05 | 7906 | 5174 | 69.08 | 4772 | 12654 |
| 10010 | 12000 | 32.29 | 1394 | 6662 | 134.81 | 7906 | 4267 | 13.11 | 4034 | 4109 |
| 10438 | 15936 | 79.68 | 1310 | 17135 | 170.90 | 7906 | 10636 | 119.29 | 5804 | 24476 |
| 10436 | 15943 | 28.69 | 1394 | 6046 | 44.49 | 7906 | 2073 | 24.36 | 4110 | 4995 |
| 10104 | 7325 | 115.72 | 1350 | 17033 | 418.09 | 7906 | 10801 | 135.72 | 5520 | 21228 |
| 10118 | 7667 | 111.69 | 1350 | 25151 | 1741.01 | 7906 | 25683 | 141.20 | 6200 | 26013 |
| 10032 | 676 | 73.29 | 1350 | 19788 | 555.74 | 7906 | 13680 | 101.64 | 5520 | 21191 |
| **AVERAGE** | | 70.81 | 1355 | 16454 | 383.54 | 7906 | 9661 | 90.27 | 5252 | 17803 |
| **STD. DEVIATION** | | 28.67 | 24.19 | 6636.66 | 509.13 | 0.00 | 6577.15 | 43.05 | 713.41 | 7791.07 |

**Supplementary Table 4:** ETSAM Stage 1 Performance across different automated prompt techniques on experimental tomograms in the test set

| CDP IDs | | Zero Mask | | | | Grid mask at only first slice | | | | Grid mask at every 50th slice | | | |
|---|---|---|---|---|---|---|---|---|---|---|---|---|---|
| **Dataset** | **Run** | **Dice/F1** | **IoU** | **Precision** | **Recall** | **Dice/F1** | **IoU** | **Precision** | **Recall** | **Dice/F1** | **IoU** | **Precision** | **Recall** |
| 10000 | 245 | 0.4942 | 0.3282 | 0.8208 | 0.3536 | 0.7423 | 0.5902 | 0.8573 | 0.6545 | 0.6274 | 0.4570 | 0.8864 | 0.4855 |
| 10000 | 247 | 0.0000 | 0.0000 | 0.0000 | 0.0000 | 0.8821 | 0.7891 | 0.8383 | 0.9308 | 0.8857 | 0.7948 | 0.9391 | 0.8381 |
| 10001 | 258 | 0.8987 | 0.8161 | 0.9177 | 0.8805 | 0.9192 | 0.8505 | 0.9131 | 0.9254 | 0.8420 | 0.7271 | 0.9666 | 0.7459 |
| 10010 | 11972 | 0.8380 | 0.7212 | 0.9642 | 0.7410 | 0.8634 | 0.7597 | 0.9045 | 0.8260 | 0.7050 | 0.5444 | 0.9876 | 0.5482 |
| 10010 | 12000 | 0.8484 | 0.7367 | 0.9002 | 0.8022 | 0.8772 | 0.7813 | 0.8899 | 0.8649 | 0.8099 | 0.6806 | 0.8904 | 0.7428 |
| 10438 | 15936 | 0.5930 | 0.4215 | 0.6490 | 0.5459 | 0.6590 | 0.4914 | 0.5937 | 0.7403 | 0.4930 | 0.3271 | 0.7305 | 0.3721 |
| 10436 | 15943 | 0.0000 | 0.0000 | 0.0000 | 0.0000 | 0.7555 | 0.6071 | 0.8710 | 0.6671 | 0.5809 | 0.4094 | 0.9092 | 0.4268 |
| 10104 | 7325 | 0.2361 | 0.1338 | 0.2188 | 0.2563 | 0.1660 | 0.0905 | 0.0947 | 0.6706 | 0.4342 | 0.2773 | 0.3903 | 0.4892 |
| 10118 | 7667 | 0.8193 | 0.6939 | 0.8327 | 0.8063 | 0.5265 | 0.3573 | 0.3878 | 0.8193 | 0.6582 | 0.4905 | 0.6675 | 0.6491 |
| 10032 | 676 | 0.0000 | 0.0000 | 0.0000 | 0.0000 | 0.7504 | 0.6005 | 0.9437 | 0.6228 | 0.5425 | 0.3722 | 0.9445 | 0.3805 |
| **AVERAGE** | | 0.4728 | 0.3851 | 0.5303 | 0.4386 | 0.7142 | 0.5918 | 0.7294 | 0.7722 | 0.6579 | 0.5080 | 0.8312 | 0.5678 |

**Supplementary Table 5:** ETSAM Stage 2 Threshold Ablation with Prompt Technique - Zero Mask | Evaluated on experimental tomograms in the test set

| CDP IDs | | > -2.0 | | | | > -1.0 | | | | > -0.5 | | | | > -0.25 | | | |
|---|---|---|---|---|---|---|---|---|---|---|---|---|---|---|---|---|---|
| Dataset | Run | Dice/F1 | IoU | Precision | Recall | Dice/F1 | IoU | Precision | Recall | Dice/F1 | IoU | Precision | Recall | Dice/F1 | IoU | Precision | Recall |
| 10000 | 245 | 0.7542 | 0.6054 | 0.7822 | 0.7282 | 0.7506 | 0.6007 | 0.8443 | 0.6756 | 0.7376 | 0.5843 | 0.8696 | 0.6404 | 0.7261 | 0.5700 | 0.8780 | 0.6191 |
| 10000 | 247 | 0.8992 | 0.8169 | 0.8288 | 0.9827 | 0.9421 | 0.8905 | 0.9090 | 0.9776 | 0.9574 | 0.9182 | 0.9474 | 0.9675 | 0.9595 | 0.9222 | 0.9623 | 0.9568 |
| 10001 | 258 | 0.8388 | 0.7223 | 0.7240 | 0.9968 | 0.9289 | 0.8672 | 0.8733 | 0.9920 | 0.9559 | 0.9156 | 0.9435 | 0.9687 | 0.9519 | 0.9082 | 0.9645 | 0.9396 |
| 10010 | 11972 | 0.6956 | 0.5333 | 0.5398 | 0.9780 | 0.8225 | 0.6985 | 0.7219 | 0.9557 | 0.8800 | 0.7858 | 0.8325 | 0.9333 | 0.8982 | 0.8152 | 0.8856 | 0.9112 |
| 10010 | 12000 | 0.8018 | 0.6692 | 0.6879 | 0.9608 | 0.8731 | 0.7748 | 0.8121 | 0.9440 | 0.9004 | 0.8189 | 0.8763 | 0.9259 | 0.9041 | 0.8250 | 0.9014 | 0.9068 |
| 10438 | 15936 | 0.5634 | 0.3922 | 0.4258 | 0.8326 | 0.6349 | 0.4651 | 0.5336 | 0.7836 | 0.6623 | 0.4951 | 0.5983 | 0.7416 | 0.6680 | 0.5015 | 0.6293 | 0.7119 |
| 10436 | 15943 | 0.6718 | 0.5058 | 0.5436 | 0.8791 | 0.7473 | 0.5966 | 0.6910 | 0.8137 | 0.7687 | 0.6243 | 0.7788 | 0.7588 | 0.7671 | 0.6222 | 0.8208 | 0.7200 |
| 10104 | 7325 | 0.2418 | 0.1375 | 0.1418 | 0.8202 | 0.3249 | 0.1940 | 0.2067 | 0.7591 | 0.3642 | 0.2227 | 0.2448 | 0.7108 | 0.3809 | 0.2352 | 0.2643 | 0.6811 |
| 10118 | 7667 | 0.7679 | 0.6232 | 0.6317 | 0.9790 | 0.8169 | 0.6904 | 0.7101 | 0.9613 | 0.8338 | 0.7150 | 0.7491 | 0.9402 | 0.8383 | 0.7216 | 0.7667 | 0.9246 |
| 10032 | 676 | 0.7951 | 0.6598 | 0.8147 | 0.7763 | 0.7909 | 0.6542 | 0.8882 | 0.7129 | 0.7805 | 0.6400 | 0.9250 | 0.6751 | 0.7726 | 0.6294 | 0.9430 | 0.6543 |
| AVERAGE | | 0.7030 | 0.5666 | 0.6120 | 0.8934 | 0.7632 | 0.6432 | 0.7190 | 0.8576 | 0.7841 | 0.6720 | 0.7765 | 0.8262 | 0.7867 | 0.6751 | 0.8016 | 0.8025 |

| CDP IDs | | > 0.0 | | | | > 0.25 | | | | > 0.5 | | | | > 1.0 | | | |
|---|---|---|---|---|---|---|---|---|---|---|---|---|---|---|---|---|---|
| Dataset | Run | Dice/F1 | IoU | Precision | Recall | Dice/F1 | IoU | Precision | Recall | Dice/F1 | IoU | Precision | Recall | Dice/F1 | IoU | Precision | Recall |
| 10000 | 245 | 0.7116 | 0.5523 | 0.8839 | 0.5955 | 0.6943 | 0.5318 | 0.8876 | 0.5702 | 0.6752 | 0.5097 | 0.8899 | 0.5440 | 0.6317 | 0.4617 | 0.8909 | 0.4894 |
| 10000 | 247 | 0.9567 | 0.9170 | 0.9730 | 0.9410 | 0.9497 | 0.9043 | 0.9800 | 0.9213 | 0.9397 | 0.8863 | 0.9844 | 0.8990 | 0.9147 | 0.8427 | 0.9895 | 0.8504 |
| 10001 | 258 | 0.9369 | 0.8812 | 0.9757 | 0.9010 | 0.9155 | 0.8442 | 0.9817 | 0.8577 | 0.8911 | 0.8036 | 0.9854 | 0.8133 | 0.8371 | 0.7199 | 0.9896 | 0.7254 |
| 10010 | 11972 | 0.8986 | 0.8158 | 0.9258 | 0.8729 | 0.8801 | 0.7859 | 0.9502 | 0.8197 | 0.8500 | 0.7391 | 0.9641 | 0.7600 | 0.7744 | 0.6319 | 0.9779 | 0.6410 |
| 10010 | 12000 | 0.8970 | 0.8133 | 0.9169 | 0.8780 | 0.8816 | 0.7883 | 0.9250 | 0.8421 | 0.8619 | 0.7574 | 0.9297 | 0.8034 | 0.8164 | 0.6898 | 0.9346 | 0.7248 |
| 10438 | 15936 | 0.6668 | 0.5002 | 0.6577 | 0.6763 | 0.6592 | 0.4916 | 0.6836 | 0.6365 | 0.6459 | 0.4770 | 0.7074 | 0.5943 | 0.6061 | 0.4348 | 0.7509 | 0.5081 |
| 10436 | 15943 | 0.7521 | 0.6027 | 0.8560 | 0.6707 | 0.7233 | 0.5665 | 0.8828 | 0.6126 | 0.6843 | 0.5202 | 0.9032 | 0.5509 | 0.5889 | 0.4174 | 0.9340 | 0.4301 |
| 10104 | 7325 | 0.3949 | 0.2460 | 0.2841 | 0.6472 | 0.4052 | 0.2541 | 0.3034 | 0.6096 | 0.4108 | 0.2585 | 0.3218 | 0.5679 | 0.4090 | 0.2570 | 0.3570 | 0.4786 |
| 10118 | 7667 | 0.8393 | 0.7231 | 0.7825 | 0.9051 | 0.8368 | 0.7195 | 0.7962 | 0.8819 | 0.8307 | 0.7104 | 0.8078 | 0.8549 | 0.8079 | 0.6777 | 0.8256 | 0.7909 |
| 10032 | 676 | 0.7618 | 0.6153 | 0.9595 | 0.6317 | 0.7473 | 0.5966 | 0.9734 | 0.6065 | 0.7283 | 0.5726 | 0.9832 | 0.5783 | 0.6788 | 0.5138 | 0.9921 | 0.5159 |
| AVERAGE | | 0.7816 | 0.6667 | 0.8215 | 0.7719 | 0.7693 | 0.6483 | 0.8364 | 0.7358 | 0.7518 | 0.6235 | 0.8477 | 0.6966 | 0.7065 | 0.5647 | 0.8642 | 0.6155 |

| CDP IDs | | > 2.0 | | | |
|---|---|---|---|---|---|
| Dataset | Run | Dice/F1 | IoU | Precision | Recall |
| 10000 | 245 | 0.5293 | 0.3599 | 0.8856 | 0.3774 |
| 10000 | 247 | 0.8534 | 0.7442 | 0.9943 | 0.7474 |
| 10001 | 258 | 0.7147 | 0.5561 | 0.9935 | 0.5581 |
| 10010 | 11972 | 0.6029 | 0.4315 | 0.9880 | 0.4338 |
| 10010 | 12000 | 0.7156 | 0.5571 | 0.9400 | 0.5777 |
| 10438 | 15936 | 0.4830 | 0.3184 | 0.8188 | 0.3425 |
| 10436 | 15943 | 0.3685 | 0.2259 | 0.9714 | 0.2274 |
| 10104 | 7325 | 0.3405 | 0.2052 | 0.4121 | 0.2901 |
| 10118 | 7667 | 0.7275 | 0.5718 | 0.8452 | 0.6386 |
| 10032 | 676 | 0.5587 | 0.3876 | 0.9975 | 0.3880 |
| AVERAGE | | 0.5894 | 0.4358 | 0.8846 | 0.4581 |

**Supplementary Table 6:** ETSAM Stage 2 Threshold Ablation with Prompt Technique - Grid mask at every 50th slice | Evaluated on experimental tomograms in the test set

| CDP IDs | | > -2.0 | | | | > -1.0 | | | | > -0.5 | | | | > -0.25 | | | |
|---|---|---|---|---|---|---|---|---|---|---|---|---|---|---|---|---|---|
| **Dataset** | **Run** | **Dice/F1** | **IoU** | **Precision** | **Recall** | **Dice/F1** | **IoU** | **Precision** | **Recall** | **Dice/F1** | **IoU** | **Precision** | **Recall** | **Dice/F1** | **IoU** | **Precision** | **Recall** |
| 10000 | 245 | 0.6386 | 0.4690 | 0.8362 | 0.5165 | 0.5672 | 0.3959 | 0.8816 | 0.4181 | 0.5183 | 0.3498 | 0.8892 | 0.3657 | 0.4914 | 0.3257 | 0.8892 | 0.3395 |
| 10000 | 247 | 0.1835 | 0.1010 | 0.6207 | 0.1076 | 0.1719 | 0.0940 | 0.7121 | 0.0978 | 0.1627 | 0.0886 | 0.7479 | 0.0913 | 0.1572 | 0.0853 | 0.7632 | 0.0877 |
| 10001 | 258 | 0.8201 | 0.6950 | 0.7871 | 0.8558 | 0.8482 | 0.7364 | 0.9416 | 0.7717 | 0.7983 | 0.6642 | 0.9696 | 0.6784 | 0.7630 | 0.6169 | 0.9748 | 0.6268 |
| 10010 | 11972 | 0.7896 | 0.6523 | 0.6989 | 0.9073 | 0.8511 | 0.7407 | 0.9054 | 0.8029 | 0.8052 | 0.6739 | 0.9640 | 0.6913 | 0.7627 | 0.6164 | 0.9758 | 0.6260 |
| 10010 | 12000 | 0.8039 | 0.6721 | 0.7689 | 0.8423 | 0.8264 | 0.7042 | 0.8959 | 0.7669 | 0.8006 | 0.6675 | 0.9342 | 0.7004 | 0.7766 | 0.6348 | 0.9436 | 0.6599 |
| 10438 | 15936 | 0.4960 | 0.3298 | 0.5747 | 0.4362 | 0.4719 | 0.3089 | 0.6863 | 0.3596 | 0.4346 | 0.2776 | 0.7303 | 0.3093 | 0.4112 | 0.2588 | 0.7492 | 0.2833 |
| 10436 | 15943 | 0.6947 | 0.5323 | 0.6654 | 0.7268 | 0.6744 | 0.5087 | 0.8348 | 0.5657 | 0.6022 | 0.4308 | 0.8924 | 0.4544 | 0.5536 | 0.3827 | 0.9119 | 0.3974 |
| 10104 | 7325 | 0.3392 | 0.2043 | 0.2208 | 0.7319 | 0.4293 | 0.2733 | 0.3257 | 0.6298 | 0.4482 | 0.2888 | 0.3764 | 0.5538 | 0.4468 | 0.2877 | 0.3982 | 0.5091 |
| 10118 | 7667 | 0.7812 | 0.6409 | 0.6971 | 0.8884 | 0.8074 | 0.6770 | 0.7917 | 0.8237 | 0.7988 | 0.6650 | 0.8315 | 0.7686 | 0.7879 | 0.6501 | 0.8480 | 0.7358 |
| 10032 | 676 | 0.6805 | 0.5157 | 0.8661 | 0.5604 | 0.6075 | 0.4363 | 0.9538 | 0.4457 | 0.5440 | 0.3736 | 0.9749 | 0.3773 | 0.5078 | 0.3403 | 0.9803 | 0.3426 |
| **AVERAGE** | | 0.6227 | 0.4812 | 0.6736 | 0.6573 | 0.6255 | 0.4875 | 0.7929 | 0.5682 | 0.5913 | 0.4480 | 0.8310 | 0.4991 | 0.5658 | 0.4199 | 0.8434 | 0.4608 |

| CDP IDs | | > 0.0 | | | | > 0.25 | | | | > 0.5 | | | | > 1.0 | | | |
|---|---|---|---|---|---|---|---|---|---|---|---|---|---|---|---|---|---|
| **Dataset** | **Run** | **Dice/F1** | **IoU** | **Precision** | **Recall** | **Dice/F1** | **IoU** | **Precision** | **Recall** | **Dice/F1** | **IoU** | **Precision** | **Recall** | **Dice/F1** | **IoU** | **Precision** | **Recall** |
| 10000 | 245 | 0.4634 | 0.3016 | 0.8869 | 0.3136 | 0.4352 | 0.2782 | 0.8836 | 0.2887 | 0.4066 | 0.2552 | 0.8787 | 0.2645 | 0.3495 | 0.2118 | 0.8655 | 0.2190 |
| 10000 | 247 | 0.1514 | 0.0819 | 0.7775 | 0.0839 | 0.1452 | 0.0783 | 0.7910 | 0.0800 | 0.1390 | 0.0747 | 0.8045 | 0.0760 | 0.1250 | 0.0667 | 0.8281 | 0.0676 |
| 10001 | 258 | 0.7246 | 0.5682 | 0.9782 | 0.5755 | 0.6841 | 0.5199 | 0.9804 | 0.5254 | 0.6424 | 0.4732 | 0.9819 | 0.4773 | 0.5572 | 0.3862 | 0.9834 | 0.3887 |
| 10010 | 11972 | 0.7142 | 0.5555 | 0.9813 | 0.5615 | 0.6631 | 0.4960 | 0.9840 | 0.5000 | 0.6113 | 0.4402 | 0.9857 | 0.4431 | 0.5096 | 0.3420 | 0.9876 | 0.3434 |
| 10010 | 12000 | 0.7483 | 0.5978 | 0.9484 | 0.6179 | 0.7180 | 0.5600 | 0.9515 | 0.5765 | 0.6862 | 0.5223 | 0.9539 | 0.5358 | 0.6219 | 0.4513 | 0.9582 | 0.4604 |
| 10438 | 15936 | 0.3859 | 0.2391 | 0.7663 | 0.2579 | 0.3592 | 0.2189 | 0.7812 | 0.2332 | 0.3318 | 0.1989 | 0.7942 | 0.2097 | 0.2771 | 0.1608 | 0.8154 | 0.1669 |
| 10436 | 15943 | 0.5015 | 0.3346 | 0.9272 | 0.3437 | 0.4476 | 0.2883 | 0.9393 | 0.2938 | 0.3942 | 0.2455 | 0.9494 | 0.2487 | 0.2928 | 0.1715 | 0.9647 | 0.1726 |
| 10104 | 7325 | 0.4385 | 0.2808 | 0.4174 | 0.4619 | 0.4239 | 0.2689 | 0.4339 | 0.4143 | 0.4032 | 0.2525 | 0.4478 | 0.3668 | 0.3489 | 0.2113 | 0.4665 | 0.2786 |
| 10118 | 7667 | 0.7723 | 0.6290 | 0.8616 | 0.6997 | 0.7527 | 0.6035 | 0.8727 | 0.6617 | 0.7289 | 0.5735 | 0.8812 | 0.6216 | 0.6727 | 0.5069 | 0.8917 | 0.5401 |
| 10032 | 676 | 0.4696 | 0.3069 | 0.9834 | 0.3085 | 0.4307 | 0.2744 | 0.9852 | 0.2756 | 0.3918 | 0.2436 | 0.9863 | 0.2444 | 0.3182 | 0.1892 | 0.9875 | 0.1896 |
| **AVERAGE** | | 0.5370 | 0.3895 | 0.8528 | 0.4224 | 0.5060 | 0.3586 | 0.8603 | 0.3849 | 0.4735 | 0.3280 | 0.8664 | 0.3488 | 0.4073 | 0.2698 | 0.8749 | 0.2827 |

| CDP IDs | | > 2.0 | | | |
|---|---|---|---|---|---|
| **Dataset** | **Run** | **Dice/F1** | **IoU** | **Precision** | **Recall** |
| 10000 | 245 | 0.2413 | 0.1372 | 0.8232 | 0.1413 |
| 10000 | 247 | 0.0950 | 0.0499 | 0.8665 | 0.0503 |
| 10001 | 258 | 0.3921 | 0.2438 | 0.9836 | 0.2448 |
| 10010 | 11972 | 0.3253 | 0.1942 | 0.9880 | 0.1947 |
| 10010 | 12000 | 0.4943 | 0.3283 | 0.9658 | 0.3322 |
| 10438 | 15936 | 0.1788 | 0.0982 | 0.8426 | 0.1000 |
| 10436 | 15943 | 0.1354 | 0.0726 | 0.9835 | 0.0727 |
| 10104 | 7325 | 0.2248 | 0.1266 | 0.4804 | 0.1467 |
| 10118 | 7667 | 0.5345 | 0.3647 | 0.8959 | 0.3809 |
| 10032 | 676 | 0.2009 | 0.1116 | 0.9888 | 0.1118 |
| **AVERAGE** | | 0.2822 | 0.1727 | 0.8818 | 0.1775 |

**Supplementary Table 7:** ETSAM Stage 2 Threshold Ablation with Prompt Technique - Grid mask at only first slice | Evaluated on experimental tomograms in the test set

| CDP IDs | | > -2.0 | | | | > -1.0 | | | | > -0.5 | | | | > -0.25 | | | |
|---|---|---|---|---|---|---|---|---|---|---|---|---|---|---|---|---|---|
| Dataset | Run | Dice/F1 | IoU | Precision | Recall | Dice/F1 | IoU | Precision | Recall | Dice/F1 | IoU | Precision | Recall | Dice/F1 | IoU | Precision | Recall |
| 10000 | 245 | 0.7543 | 0.6056 | 0.7622 | 0.7466 | 0.7555 | 0.6070 | 0.8298 | 0.6933 | 0.7476 | 0.5969 | 0.8612 | 0.6604 | 0.7387 | 0.5857 | 0.8730 | 0.6402 |
| 10000 | 247 | 0.8899 | 0.8017 | 0.8137 | 0.9818 | 0.9332 | 0.8747 | 0.8942 | 0.9756 | 0.9509 | 0.9063 | 0.9351 | 0.9672 | 0.9559 | 0.9155 | 0.9531 | 0.9587 |
| 10001 | 258 | 0.8267 | 0.7046 | 0.7058 | 0.9976 | 0.9200 | 0.8518 | 0.8560 | 0.9942 | 0.9572 | 0.9179 | 0.9352 | 0.9802 | 0.9581 | 0.9196 | 0.9609 | 0.9554 |
| 10010 | 11972 | 0.7150 | 0.5564 | 0.5634 | 0.9781 | 0.8317 | 0.7119 | 0.7354 | 0.9571 | 0.8867 | 0.7965 | 0.8412 | 0.9375 | 0.9058 | 0.8278 | 0.8933 | 0.9186 |
| 10010 | 12000 | 0.7956 | 0.6606 | 0.6781 | 0.9623 | 0.8699 | 0.7698 | 0.8057 | 0.9453 | 0.9003 | 0.8187 | 0.8736 | 0.9287 | 0.9052 | 0.8269 | 0.9003 | 0.9102 |
| 10438 | 15936 | 0.5477 | 0.3771 | 0.3963 | 0.8864 | 0.6266 | 0.4562 | 0.4991 | 0.8416 | 0.6633 | 0.4962 | 0.5633 | 0.8063 | 0.6758 | 0.5104 | 0.5957 | 0.7810 |
| 10436 | 15943 | 0.6692 | 0.5029 | 0.5389 | 0.8826 | 0.7442 | 0.5926 | 0.6846 | 0.8152 | 0.7666 | 0.6216 | 0.7721 | 0.7612 | 0.7672 | 0.6223 | 0.8154 | 0.7243 |
| 10104 | 7325 | 0.1589 | 0.0863 | 0.0880 | 0.8176 | 0.1939 | 0.1073 | 0.1110 | 0.7657 | 0.2092 | 0.1168 | 0.1223 | 0.7240 | 0.2153 | 0.1207 | 0.1274 | 0.6968 |
| 10118 | 7667 | 0.7538 | 0.6048 | 0.6117 | 0.9817 | 0.8072 | 0.6767 | 0.6929 | 0.9667 | 0.8282 | 0.7068 | 0.7347 | 0.9490 | 0.8353 | 0.7171 | 0.7544 | 0.9356 |
| 10032 | 676 | 0.7936 | 0.6578 | 0.8032 | 0.7841 | 0.7902 | 0.6532 | 0.8805 | 0.7168 | 0.7796 | 0.6389 | 0.9195 | 0.6767 | 0.7714 | 0.6279 | 0.9383 | 0.6549 |
| AVERAGE | | 0.6905 | 0.5558 | 0.5961 | 0.9019 | 0.7472 | 0.6301 | 0.6989 | 0.8672 | 0.7690 | 0.6617 | 0.7558 | 0.8391 | 0.7729 | 0.6674 | 0.7812 | 0.8176 |

| CDP IDs | | > 0.0 | | | | > 0.25 | | | | > 0.5 | | | | > 1.0 | | | |
|---|---|---|---|---|---|---|---|---|---|---|---|---|---|---|---|---|---|
| Dataset | Run | Dice/F1 | IoU | Precision | Recall | Dice/F1 | IoU | Precision | Recall | Dice/F1 | IoU | Precision | Recall | Dice/F1 | IoU | Precision | Recall |
| 10000 | 245 | 0.7258 | 0.5696 | 0.8815 | 0.6169 | 0.7095 | 0.5498 | 0.8871 | 0.5911 | 0.6908 | 0.5276 | 0.8906 | 0.5642 | 0.6479 | 0.4791 | 0.8940 | 0.5080 |
| 10000 | 247 | 0.9564 | 0.9165 | 0.9672 | 0.9459 | 0.9520 | 0.9085 | 0.9772 | 0.9281 | 0.9435 | 0.8930 | 0.9836 | 0.9065 | 0.9189 | 0.8500 | 0.9900 | 0.8573 |
| 10001 | 258 | 0.9447 | 0.8952 | 0.9736 | 0.9174 | 0.9237 | 0.8583 | 0.9797 | 0.8738 | 0.8994 | 0.8172 | 0.9832 | 0.8288 | 0.8463 | 0.7335 | 0.9869 | 0.7407 |
| 10010 | 11972 | 0.9074 | 0.8305 | 0.9325 | 0.8836 | 0.8897 | 0.8013 | 0.9550 | 0.8327 | 0.8606 | 0.7554 | 0.9669 | 0.7754 | 0.7892 | 0.6518 | 0.9779 | 0.6615 |
| 10010 | 12000 | 0.8979 | 0.8146 | 0.9163 | 0.8801 | 0.8819 | 0.7887 | 0.9242 | 0.8432 | 0.8614 | 0.7565 | 0.9284 | 0.8034 | 0.8150 | 0.6878 | 0.9327 | 0.7238 |
| 10438 | 15936 | 0.6817 | 0.5171 | 0.6255 | 0.7489 | 0.6802 | 0.5153 | 0.6523 | 0.7105 | 0.6723 | 0.5063 | 0.6764 | 0.6682 | 0.6419 | 0.4727 | 0.7200 | 0.5791 |
| 10436 | 15943 | 0.7551 | 0.6066 | 0.8529 | 0.6774 | 0.7287 | 0.5731 | 0.8805 | 0.6215 | 0.6918 | 0.5288 | 0.9010 | 0.5615 | 0.6011 | 0.4297 | 0.9309 | 0.4439 |
| 10104 | 7325 | 0.2202 | 0.1237 | 0.1319 | 0.6649 | 0.2233 | 0.1257 | 0.1358 | 0.6277 | 0.2250 | 0.1268 | 0.1392 | 0.5865 | 0.2252 | 0.1269 | 0.1455 | 0.4975 |
| 10118 | 7667 | 0.8394 | 0.7233 | 0.7727 | 0.9187 | 0.8401 | 0.7243 | 0.7892 | 0.8981 | 0.8375 | 0.7204 | 0.8040 | 0.8739 | 0.8211 | 0.6965 | 0.8276 | 0.8147 |
| 10032 | 676 | 0.7603 | 0.6133 | 0.9561 | 0.6311 | 0.7455 | 0.5943 | 0.9711 | 0.6050 | 0.7258 | 0.5696 | 0.9818 | 0.5757 | 0.6746 | 0.5090 | 0.9918 | 0.5111 |
| AVERAGE | | 0.7689 | 0.6610 | 0.8010 | 0.7885 | 0.7575 | 0.6439 | 0.8152 | 0.7532 | 0.7408 | 0.6202 | 0.8255 | 0.7144 | 0.6981 | 0.5637 | 0.8397 | 0.6338 |

| CDP IDs | | > 2.0 | | | |
|---|---|---|---|---|---|
| Dataset | Run | Dice/F1 | IoU | Precision | Recall |
| 10000 | 245 | 0.5457 | 0.3752 | 0.8914 | 0.3932 |
| 10000 | 247 | 0.8559 | 0.7481 | 0.9949 | 0.7510 |
| 10001 | 258 | 0.7281 | 0.5724 | 0.9902 | 0.5757 |
| 10010 | 11972 | 0.6317 | 0.4616 | 0.9852 | 0.4648 |
| 10010 | 12000 | 0.7148 | 0.5562 | 0.9373 | 0.5777 |
| 10438 | 15936 | 0.5354 | 0.3655 | 0.7899 | 0.4049 |
| 10436 | 15943 | 0.3929 | 0.2445 | 0.9687 | 0.2464 |
| 10104 | 7325 | 0.2106 | 0.1177 | 0.1589 | 0.3122 |
| 10118 | 7667 | 0.7511 | 0.6014 | 0.8562 | 0.6689 |
| 10032 | 676 | 0.5520 | 0.3812 | 0.9981 | 0.3815 |
| AVERAGE | | 0.5918 | 0.4424 | 0.8571 | 0.4776 |

**Supplementary Table 8:** ETSAM Stage 2 Threshold Ablation with Prompt Technique - Stage 1 mask at every 50th slice | Evaluated on experimental tomograms in the test set

| CDP IDs | | > -2.0 | | | | > -1.0 | | | | > -0.5 | | | | > -0.25 | | | |
|---|---|---|---|---|---|---|---|---|---|---|---|---|---|---|---|---|---|
| Dataset | Run | Dice/F1 | IoU | Precision | Recall | Dice/F1 | IoU | Precision | Recall | Dice/F1 | IoU | Precision | Recall | Dice/F1 | IoU | Precision | Recall |
| 10000 | 245 | 0.7743 | 0.6318 | 0.8133 | 0.7390 | 0.7701 | 0.6261 | 0.8711 | 0.6900 | 0.7629 | 0.6167 | 0.9004 | 0.6619 | 0.7486 | 0.5982 | 0.9038 | 0.6388 |
| 10000 | 247 | 0.9068 | 0.8295 | 0.8435 | 0.9803 | 0.9481 | 0.9012 | 0.9221 | 0.9755 | 0.9608 | 0.9245 | 0.9580 | 0.9635 | 0.9517 | 0.9078 | 0.9616 | 0.9420 |
| 10001 | 258 | 0.8564 | 0.7488 | 0.7498 | 0.9983 | 0.9398 | 0.8864 | 0.8883 | 0.9975 | 0.9812 | 0.9631 | 0.9688 | 0.9939 | 0.9639 | 0.9304 | 0.9744 | 0.9537 |
| 10010 | 11972 | 0.8087 | 0.6788 | 0.6936 | 0.9696 | 0.9030 | 0.8231 | 0.8657 | 0.9436 | 0.9432 | 0.8925 | 0.9657 | 0.9217 | 0.9159 | 0.8448 | 0.9717 | 0.8661 |
| 10010 | 12000 | 0.8322 | 0.7126 | 0.7369 | 0.9559 | 0.8955 | 0.8108 | 0.8554 | 0.9396 | 0.9225 | 0.8562 | 0.9191 | 0.9259 | 0.9066 | 0.8292 | 0.9238 | 0.8900 |
| 10438 | 15936 | 0.5924 | 0.4209 | 0.4518 | 0.8602 | 0.6562 | 0.4883 | 0.5480 | 0.8177 | 0.6853 | 0.5213 | 0.6056 | 0.7891 | 0.6829 | 0.5185 | 0.6229 | 0.7558 |
| 10436 | 15943 | 0.7309 | 0.5759 | 0.6525 | 0.8307 | 0.7735 | 0.6307 | 0.7898 | 0.7579 | 0.7782 | 0.6369 | 0.8693 | 0.7043 | 0.7504 | 0.6006 | 0.8837 | 0.6521 |
| 10104 | 7325 | 0.1985 | 0.1102 | 0.1131 | 0.8140 | 0.2414 | 0.1373 | 0.1435 | 0.7589 | 0.2571 | 0.1475 | 0.1569 | 0.7105 | 0.2616 | 0.1505 | 0.1620 | 0.6784 |
| 10118 | 7667 | 0.5564 | 0.3854 | 0.3900 | 0.9700 | 0.6033 | 0.4320 | 0.4451 | 0.9362 | 0.6175 | 0.4466 | 0.4696 | 0.9013 | 0.6211 | 0.4505 | 0.4804 | 0.8786 |
| 10032 | 676 | 0.7994 | 0.6658 | 0.8548 | 0.7507 | 0.7868 | 0.6486 | 0.9232 | 0.6856 | 0.7729 | 0.6298 | 0.9568 | 0.6483 | 0.7637 | 0.6177 | 0.9736 | 0.6282 |
| **AVERAGE** | | 0.7056 | 0.5760 | 0.6299 | 0.8869 | 0.7518 | 0.6385 | 0.7252 | 0.8503 | 0.7682 | 0.6635 | 0.7770 | 0.8220 | 0.7566 | 0.6448 | 0.7858 | 0.7884 |

| CDP IDs | | > 0.0 | | | | > 0.25 | | | | > 0.5 | | | | > 1.0 | | | |
|---|---|---|---|---|---|---|---|---|---|---|---|---|---|---|---|---|---|
| Dataset | Run | Dice/F1 | IoU | Precision | Recall | Dice/F1 | IoU | Precision | Recall | Dice/F1 | IoU | Precision | Recall | Dice/F1 | IoU | Precision | Recall |
| 10000 | 245 | 0.7322 | 0.5776 | 0.9055 | 0.6147 | 0.7149 | 0.5563 | 0.9068 | 0.5901 | 0.6967 | 0.5346 | 0.9076 | 0.5653 | 0.6576 | 0.4899 | 0.9083 | 0.5154 |
| 10000 | 247 | 0.9404 | 0.8874 | 0.9632 | 0.9186 | 0.9279 | 0.8655 | 0.9641 | 0.8944 | 0.9147 | 0.8428 | 0.9645 | 0.8698 | 0.8860 | 0.7953 | 0.9648 | 0.8191 |
| 10001 | 258 | 0.9431 | 0.8923 | 0.9770 | 0.9114 | 0.9211 | 0.8538 | 0.9793 | 0.8695 | 0.8983 | 0.8154 | 0.9813 | 0.8282 | 0.8498 | 0.7388 | 0.9844 | 0.7476 |
| 10010 | 11972 | 0.8848 | 0.7934 | 0.9744 | 0.8103 | 0.8528 | 0.7433 | 0.9765 | 0.7569 | 0.8197 | 0.6945 | 0.9781 | 0.7055 | 0.7523 | 0.6030 | 0.9802 | 0.6104 |
| 10010 | 12000 | 0.8880 | 0.7986 | 0.9261 | 0.8530 | 0.8685 | 0.7675 | 0.9279 | 0.8162 | 0.8482 | 0.7365 | 0.9295 | 0.7800 | 0.8060 | 0.6750 | 0.9323 | 0.7098 |
| 10438 | 15936 | 0.6775 | 0.5123 | 0.6391 | 0.7207 | 0.6699 | 0.5037 | 0.6556 | 0.6850 | 0.6602 | 0.4927 | 0.6721 | 0.6487 | 0.6333 | 0.4634 | 0.7053 | 0.5747 |
| 10436 | 15943 | 0.7183 | 0.5604 | 0.8967 | 0.5990 | 0.6825 | 0.5180 | 0.9095 | 0.5462 | 0.6436 | 0.4744 | 0.9216 | 0.4944 | 0.5577 | 0.3867 | 0.9424 | 0.3960 |
| 10104 | 7325 | 0.2640 | 0.1521 | 0.1661 | 0.6428 | 0.2644 | 0.1523 | 0.1692 | 0.6042 | 0.2626 | 0.1512 | 0.1713 | 0.5629 | 0.2545 | 0.1458 | 0.1735 | 0.4775 |
| 10118 | 7667 | 0.6223 | 0.4517 | 0.4901 | 0.8522 | 0.6212 | 0.4505 | 0.4990 | 0.8227 | 0.6178 | 0.4470 | 0.5071 | 0.7904 | 0.6048 | 0.4335 | 0.5216 | 0.7196 |
| 10032 | 676 | 0.7521 | 0.6027 | 0.9889 | 0.6068 | 0.7298 | 0.5745 | 0.9920 | 0.5772 | 0.7053 | 0.5448 | 0.9933 | 0.5468 | 0.6538 | 0.4857 | 0.9956 | 0.4867 |
| **AVERAGE** | | 0.7423 | 0.6229 | 0.7927 | 0.7530 | 0.7253 | 0.5985 | 0.7980 | 0.7162 | 0.7067 | 0.5734 | 0.8026 | 0.6792 | 0.6656 | 0.5217 | 0.8108 | 0.6057 |

| CDP IDs | | > 2.0 | | | |
|---|---|---|---|---|---|
| Dataset | Run | Dice/F1 | IoU | Precision | Recall |
| 10000 | 245 | 0.5688 | 0.3974 | 0.9055 | 0.4146 |
| 10000 | 247 | 0.8213 | 0.6968 | 0.9646 | 0.7150 |
| 10001 | 258 | 0.7430 | 0.5911 | 0.9877 | 0.5955 |
| 10010 | 11972 | 0.6123 | 0.4412 | 0.9817 | 0.4449 |
| 10010 | 12000 | 0.7151 | 0.5566 | 0.9371 | 0.5781 |
| 10438 | 15936 | 0.5480 | 0.3774 | 0.7663 | 0.4265 |
| 10436 | 15943 | 0.3745 | 0.2304 | 0.9718 | 0.2319 |
| 10104 | 7325 | 0.2185 | 0.1226 | 0.1689 | 0.3094 |
| 10118 | 7667 | 0.5569 | 0.3859 | 0.5478 | 0.5664 |
| 10032 | 676 | 0.5389 | 0.3689 | 0.9987 | 0.3691 |
| **AVERAGE** | | 0.5697 | 0.4168 | 0.8230 | 0.4651 |

**Supplementary Table 9:** Per-Tomogram Performance of ETSAM (single stage) on experimental tomograms in the test set

| CDP IDs | | ETSAM | | | |
|---|---|---|---|---|---|
| **Dataset** | **Run** | **Dice/F1** | **IoU** | **Precision** | **Recall** |
| 10000 | 245 | 0.7423 | 0.5902 | 0.8573 | 0.6545 |
| 10000 | 247 | 0.8821 | 0.7891 | 0.8383 | 0.9308 |
| 10001 | 258 | 0.9192 | 0.8505 | 0.9131 | 0.9254 |
| 10010 | 11972 | 0.8634 | 0.7597 | 0.9045 | 0.8260 |
| 10010 | 12000 | 0.8772 | 0.7813 | 0.8899 | 0.8649 |
| 10438 | 15936 | 0.6590 | 0.4914 | 0.5937 | 0.7403 |
| 10436 | 15943 | 0.7555 | 0.6071 | 0.8710 | 0.6671 |
| 10104 | 7325 | 0.1660 | 0.0905 | 0.0947 | 0.6706 |
| 10118 | 7667 | 0.5265 | 0.3573 | 0.3878 | 0.8193 |
| 10032 | 676 | 0.7504 | 0.6005 | 0.9437 | 0.6228 |
| **AVERAGE** | | 0.7142 | 0.5918 | 0.7294 | 0.7722 |
